# FISIK: Framework for the Inference of in Situ Interaction Kinetics from single-molecule imaging data

**DOI:** 10.1101/530063

**Authors:** L. R. de Oliveira, K. Jaqaman

## Abstract

Recent experimental and computational developments have been pushing the limits of live-cell single-molecule imaging, enabling the monitoring of inter-molecular interactions in their native environment with high spatiotemporal resolution. However, interactions are captured only for the labeled subset of molecules, which tends to be a small fraction. As a result, it has remained a challenge to calculate molecular interaction kinetics, in particular association rates, from live-cell single-molecule tracking data. To overcome this challenge, we developed a mathematical modeling-based Framework for the Inference of in Situ Interaction Kinetics from single-molecule imaging data with sub-stoichiometric labeling (termed “FISIK”). FISIK consists of (I) devising a mathematical model of molecular movement and interactions, mimicking the biological system and data-acquisition setup, and (II) estimating the unknown model parameters, including molecular association and dissociation rates, by fitting the model to experimental single-molecule data. Due to the stochastic nature of the model and data, we adapted the method of indirect inference for model calibration. We validated FISIK using a series of tests, where we simulated trajectories of diffusing molecules that interact with each other, considering a wide range of model parameters, and including resolution limitations, tracking errors and mismatches between the model and the biological system it mimics. We found that FISIK has the sensitivity to determine association and dissociation rates, with accuracy and precision depending on the labeled fraction of molecules and the extent of molecule tracking errors. For cases where the labeled fraction is too low (e.g. to afford accurate tracking), combining dynamic but sparse single-molecule imaging data with almost whole-population oligomer distribution data improves FISIK’s performance. All in all, FISIK is a promising approach for the derivation of molecular interaction kinetics in their native environment from single-molecule imaging data with sub-stoichiometric labeling.

**Significance:** Live-cell single-molecule imaging has the unique power to capture inter-molecular interactions in their native environment. However, single-molecule approaches in cells suffer from the inherent limitation that only a small fraction of molecules can be visualized at a time. Therefore, it has remained a challenge to calculate interaction rates, especially association rates, from these data. We have developed a mathematical modeling and model calibration-based framework (FISIK) to address this challenge, and derive molecular interaction rates from the subset of interactions captured by single-molecule imaging with sub-stoichiometric labeling. FISIK is a general framework, not limited to any particular interaction model, and is thus expected to be widely applicable, allowing the full use of the rich information provided by single-molecule imaging experiments.

## Introduction

Light microscopy is providing information on molecular activities in cells with ever-increasing sensitivity and spatial and temporal resolution. Modalities such as live-cell single-molecule imaging provide the highest sensitivity and resolution, reporting on the dynamic activities of individual molecules. Recent experimental and computational developments have been pushing the limits of single-molecule imaging to capture not only molecular movement – the traditional strength of single-molecule imaging (1-3) – but also inter-molecular interactions (4-9). Capturing molecular interactions in their native cellular environment provides critical spatial and temporal context for the observed interactions. For example, receptor interactions in their native plasma membrane environment can have different kinetics from the interactions observed via traditional 3D in vitro methods (5, 10, 11). These kinetics might also depend on the cellular environment and subcellular context (12). Additionally, microscopy approaches allow the study of interactions between proteins, such as transmembrane proteins, that are difficult to purify for traditional biophysical studies.

However, most live-cell single-molecule approaches, from direct one-color and multi-color imaging (4, 6-8, 13, 14) to single-molecule Förster resonance energy transfer (smFRET) (5, 15), suffer from the inherent limitation that only a small fraction of molecules can be visualized at a time. Such sub-stoichiometric labeling is necessary because of diffraction, which limits resolution to ∼200 nm laterally and ∼500 nm axially, and because of the need to track the imaged molecules over time (16). As a consequence of sub-stoichiometric labeling, it is challenging to calculate molecular interaction rates from single-molecule experiments, in spite of the rich information they might provide. This is because only a small subset of interaction events can be captured in single-molecule imaging experiments. Importantly, this subset is biased toward oligomers/complexes/clusters with a smaller number of molecules. For example, if 10% of molecules are labeled, then only 1% of dimers would be fully labeled and thus visualized as dimers, 0.1% of trimers, etc.

To the best of our knowledge, there are less than a handful of previous studies that have estimated molecular interaction rates from single-molecule data with sub-stoichiometric labeling. These studies were primarily focused on the special cases of dimerization (8, 17) and bimolecular interactions (14). Additionally, to derive association rates, they employed single-molecule experiments with a high-labeled fraction, greater than 75% (14, 17). Such a high degree of labeling is generally not compatible with single-molecule imaging for most molecules at their endogenous expression levels.

To fill this technological void, here we present a generic, mathematical modeling-based approach to calculate interaction kinetics from single-molecule data with sub-stoichiometric labeling, termed FISIK (Framework for the Inference of in Situ Interaction Kinetics). FISIK consists of two components: (I) devising a mathematical model that mimics the dynamics, interactions and sub-stoichiometric labeling of the molecular system, and (II) calibrating the model with single-molecule data to estimate the unknown model parameters, including the molecular association and dissociation rates. To test the feasibility, strengths and limitations of FISIK, here we applied it to the case of movements and interactions in 2D, applicable to e.g. cell-surface receptors, a system often studied with single-molecule imaging (4-9). To the best of our knowledge, this is the first attempt to develop a generic framework to derive molecular interaction rates from single-molecule data with sub-stoichiometric labeling, not limited to dimerization/bimolecular interactions. Our proof-of-principle tests also shed light on experimental design strategies that can be employed to maximize the performance of FISIK and derive the molecular interaction rates of interest.

## Methods

### 1. Overall workflow of FISIK

#### FISIK consists of two components (Fig 1)

##### Component I. A stochastic mathematical model of molecular movement and interactions, mimicking the biological system and data acquisition setup, including sparse labeling

Many parameters in this model are derived explicitly from the experimental data and directly input into the model, such as molecular movement properties and most data acquisition parameters (e.g. sampling rate). This constrains the modeling and leaves only a few unknown model parameters, including the rates of molecular association and dissociation, to be estimated by fitting the model to experimental single-molecule data. The specific model employed to test the easibility of FISIK is described in subsection 2 below.

**Figure 1.**
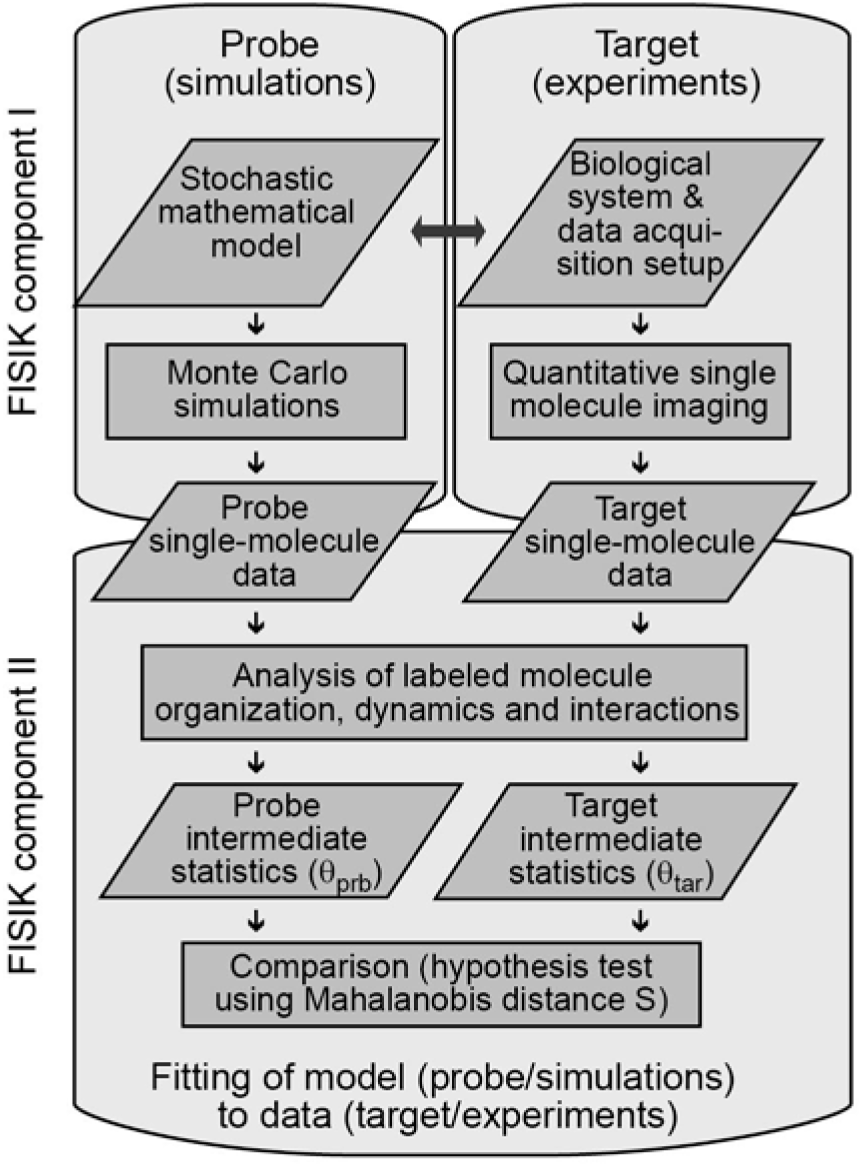
FISIK workflow. FISIK consists of two components: (I) Building a mathematical model and generating simulated data (probe) that mimic the biological system and acquired experimental data (target). (II) Fitting the model to target data in order to determine the unknown model parameters, including molecular association and dissociation rates. This requires a stochastic model calibration framework, as both model and data are stochastic.

##### Component II. Model Calibration with experimental single-molecule data to estimate molecular association and dissociation rates

The stochasticity of molecular movement and interactions, compounded by the labeling of a random subset of molecules, makes it meaningless to compare model-generated and experimental molecular trajectories and interaction instances point-by-point for model calibration, as would be done for a deterministic system. Therefore, we have devised a stochastic model calibration algorithm by adapting the method of Indirect Inference (18-20). In this approach, the model-generated (termed “probe”) and experimental (termed “target”) data are first analyzed to extract from them descriptors, called intermediate statistics, which characterize them. The specific intermediate statistics used for calibrating our test model, which are closely related to the unknown model parameters, are discussed in subsection 4 below. The choice of intermediate statistics is critical for the success of statistical inference methods such as Indirect Inference. The intermediate statistics should be sufficient, i.e. they should capture the important information about the system, yet they should minimize redundancy (21, 22). The difference between probe and target is then taken as the difference between their respective intermediate statistics, calculated as the Mahalanobis distance between them (23). Specifically, if *θ*_*prb*_ and *θ*_*tar*_ are, respectively, the intermediate statistics of the probe and target data, with associated variance-covariance matrices *V*_*prb*_ and *V*_*tar*_, then the Mahalanobis distance *S* is calculated as:

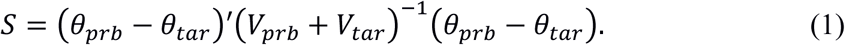

Casting the comparison in terms of the Mahalanobis distance has two advantages: (i) The difference between intermediate statistics gets weighted by their variance-covariance matrices, thus accounting for the statistics’ uncertainties when calculating the difference between target and probe. (ii) Under the null hypothesis that *θ*_*prb*_ = *θ*_*tar*_, and given that the intermediate statistics are normally distributed (as verified using a Lilliefors test, which gave p-values in the range 0.12-0.5), S follows a *χ*^2^-distribution with number of degrees of freedom equal to number of intermediate statistics. This allows us to calculate a comparison p-value for any given *S*, facilitating the interpretation of the value of *S* (24, 25). Specifically, probes with comparison p-value < α (e.g. 0.05) are considered statistically different from the target, and are thus outside the solution range. Probes with p-value ≥ α are statistically indistinguishable from the target, and are thus possible solutions.

Note that, at the conceptual level, FISIK is not limited to using Indirect Inference for stochastic model calibration. Other methods, such as Bayesian Inference (26, 27), including Approximate Bayesian Computation (28-30), are expected to be equally applicable.

### 2. Stochastic model of molecular movement, interactions and sub-stoichiometric labeling

To test the feasibility of FISIK, we employed a simple stochastic model of molecular diffusion and interactions in 2D, as well as sub-stoichiometric labeling to mimic single-molecule sampling. A 2D model is less computationally intensive to simulate than a 3D model, while at the same time providing equal power to test the feasibility of FISIK. Additionally, a 2D model is a reasonable approximation of e.g. receptor movement on the cell surface, a system that is particularly accessible and heavily studied by single-molecule imaging approaches (4-9). Membrane proteins are also challenging to purify for traditional in vitro biophysical methods, making cell-based imaging approaches particularly useful to study them. The model contains only one molecular species (e.g. a particular receptor type), which can oligomerize into dimers, trimers, etc. Note that in the following and subsequent descriptions, oligomer(*n*) indicates an oligomer of size *n*, where *n* = 1 means monomer, *n* = 2 means dimer, *n* = 3 means trimer, etc. The terms monomer and oligomer(1) will be used interchangeably, as needed for compactness of description.

In this simple model, molecules have a 2D density *ρ* and undergo 2D free diffusion with diffusion coefficient *D*. When a monomer and an oligomer(*n* ≥ 1) encounter each other, i.e. their pairwise distance ≤ *d*_size_ (taken as 10 nm, assuming a molecular radius of ∼ 5 nm), they associate with probability *p*_*a*_(*n*+1) to form an oligomer(*n*+1). The case of *n* = 1 is the special case of two monomers associating to form a dimer. Molecules in an oligomer diffuse together, with the same diffusion coefficient *D*. For an oligomer(*n* ≥ 2), a molecule can dissociate from it with rate *k*_*off*_(*n*), to produce a monomer and an oligomer(*n*-1) (note that association and dissociation are limited to one molecule at a time).

To mimic single-molecule imaging with sub-stoichiometric labeling, a fraction *f* of the molecules is labeled, with individual fluorophore intensity *I*_*ind*_ ∼ *N*(*μ*_*ind*_,*σ*_*ind*_) (for *σ*_*ind*_ > 0, the intensity of an individual fluorophore fluctuates over time, mimicking realistic fluorophore intensity fluctuations). The remaining fraction of molecules (1-*f*) is invisible. Therefore, while all molecules diffuse and interact, only the trajectories of labeled molecules and the interactions between labeled molecules are visible. Note that if a labeled molecule interacts with an unlabeled molecule, that interaction is not visible and is not recorded in the output trajectory of the labeled molecule.

The unknown parameters in this model are the motion parameter *D*, the interaction parameters *p*_*a*_(*n*) and *k*_*off*_(*n*) (*n* = 2, 3, 4, …), the population parameters *ρ* and *f*, and the fluorophore intensity parameters *μ*_*ind*_ and *σ*_*ind*_. Among these parameters, *D* can be estimated directly from the single-molecule data, as it is not affected by under-sampling. The fluorophore intensity parameters are simplified to *μ*_*ind*_ = 1 and *σ*_*ind*_ = 0 for probe simulations, but can otherwise be estimated experimentally as well, using e.g. monodispersion experiments (6). The remaining parameters, however, must be estimated by fitting the model to experimental data, as described in the general framework above (subsection 1), and using the intermediate statistics described in subsection 4 below.

### 3. Kinetic Monte Carlo simulations

To generate molecular trajectories and interactions from the above model, we used kinetic Monte Carlo simulations explicit in both space and time (effectively particle-based reaction-diffusion simulations, similar to (31, 32), although in 2D instead of 3D). In these simulations, space is treated as a continuum, while time is discretized into a series of time steps *Δt*, where *Δt* is small enough to minimize discretization artifacts (more on this below). To simulate molecules at density *ρ* within a simulation area *A, N*_*mol*_ = *ρ* × *A* molecules are initially placed randomly within the simulation area, all starting as monomers. Reflecting boundary conditions are used to keep the molecules within the simulation area. Each simulation starts with an initialization time *T*_*init*_ (usually 10 s) so that the system reaches steady state (Fig S1A; Table S1, Row 1), followed by the desired simulation time *T*_*sim*_. Only the post-initialization time (i.e. steady-state) trajectories and interactions are output for further processing and analysis.

At every time point *t* after the initial time point, molecules can dissociate, move and associate, in this order, as described in the following:

#### (1) Molecule dissociation

For each oligomer(*n* ≥ 2) at the previous time point (*t* – Δ*t*), one molecule can dissociate from it at time point *t* with probability *p*_*off*_(*n*) = *k*_*off*_(*n*) × Δ*t*, producing a monomer and an oligomer(*n*-1). If a dissociation happens, the involved monomer and oligomer are not allowed to associate with each other or with any other monomer or oligomer in the current time point (step 3 below), to avoid what will appear as instantaneous swapping of molecules between oligomers in one time point. This constraint is acceptable as long as the time step is small enough, as demonstrated by our validation tests (Fig 2).

**Figure 2.**
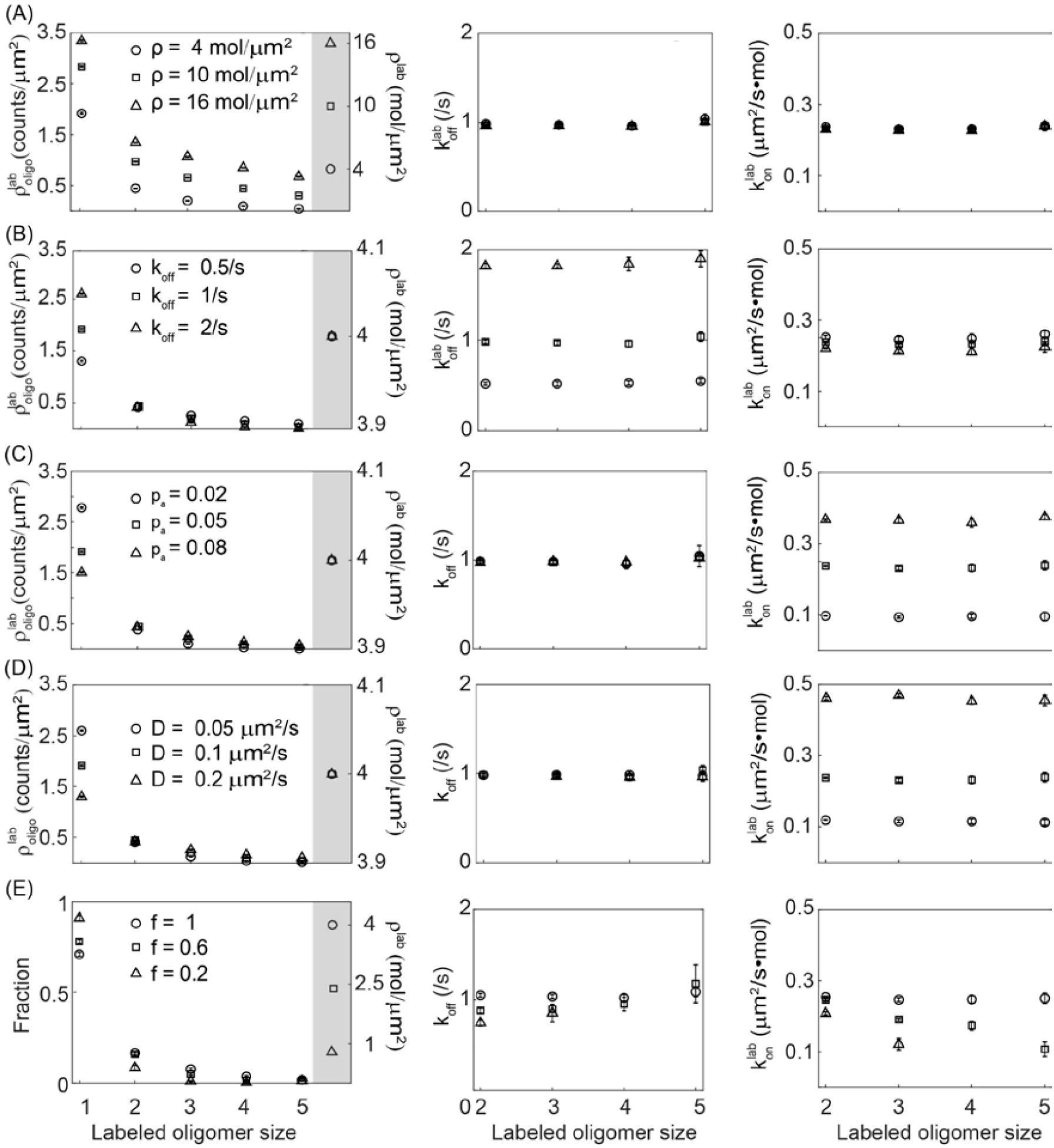
The intermediate statistics accurately reflect the modeled interactions. (**A-E**) Influence of varying molecule density *ρ* (**A**), dissociation rate *k*_*off*_ (**B**), association probability *p*_*a*_ (**C**), diffusion coefficient D (**D**) and labeled fraction *f* (**E**) on the intermediate statistics 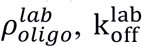 and 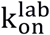. Shown are the mean and standard deviation from 10 simulations per parameter combination. Each panel explicitly states the varied parameter values, while the parameters not stated take on the following values: *ρ* = 4 mol/µm^2^, *p*_*a*_ = 0.05, *k*_*off*_ = 1/s, *D* = 0.1 µm^2^/s and *f* = 1. In the first column, the right-most measurement (with gray background) is the total molecule density (*ρ*^*lab*^) as calculated from the oligomer densities: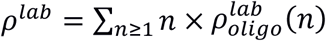. Note that the left and right y-axes have different ranges. (E) shows the oligomer fraction instead of oligomer density to illustrate better the shift toward smaller labeled oligomer sizes as the labeled fraction decreases.

#### (2) Initial update of molecule positions

Under the model of free diffusion, each oligomer(*n* ≥ 1) takes a step from time point *t* – Δ*t* to time point *t* with x- and y-components, *s*_*x*_ and 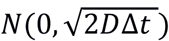, where *D* is the diffusion coefficient. As mentioned above, all molecules within an oligomer move together. Note that if a molecule has dissociated from an oligomer (step 1 above), it moves independently of the oligomer to which it used to belong. The resulting positions are considered “initial positions” because, as described next, these positions might be altered by association.

#### (3) Molecule association and final update of positions

From the model definition, if the distance between two molecules ≤ *d*_size_ (= 10 nm) at time point *t*, they are considered to encounter each other and thus might associate. However, due to time discretization in the simulations, molecules might pass by each other as they move from time point *t* – Δ*t* to time point *t*, but the distance between them might be greater than *d*_size_ at both time points. Therefore, we devised a simulation strategy to compensate for the effect of finite Δ*t* on molecular encounters. The compensation strategy is based on assigning molecules an “effective radius” (*R*_*eff*_) based on their extent of movement within Δ*t* rather than their physical size, defined as 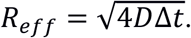 This allows us to define a movement-based encounter distance, *d*_move_ = 2*R*_*eff*_. For example, for *D* = 0.1 μm^2^/s, *d*_move_ = 12.6 nm (≈ *d*_size_ = 10 nm) for *Δt* = 10^-4^ s, but *d*_move_ = 126 nm for *Δt* = 10^-2^ s. In the simulation, the encounter distance *d*_encounter_ is then taken as max(*d*_size_, *d*_move_). Testing this compensation strategy demonstrated that it renders the simulation output insensitive to Δ*t* up to 10^-2^s (Fig S1B), a practically feasible Δ*t*, which is more than 2 orders of magnitude larger than the Δ*t* needed to avoid time discretization artifacts.

With this, all monomer-oligomer(*n* ≥ 1) pairs with pairwise distance ≤ *d*_encounter_ are considered as encounter candidates. If any objects (monomers or oligomers) appear in more than one pair because their distance ≤ *d*_encounter_ to more than one object, graph matching is used to resolve these conflicts globally (33) and impose that each object may interact with only one other object per time point. With this, monomer-oligomer(*n* ≥ 1) pairs that encounter each other can associate with probability *p*_*a*_(*n*+1) to form an oligomer(*n*+1). When an association happens, the position of the resulting oligomer is the average position of the two associating objects. This yields the final positions of molecules at time *t* of the simulation.

##### Particle tracks output of kinetic Monte Carlo simulations

The trajectories output by the simulation code to use for further analysis within FISIK are in the same form as tracks obtained via multiple particle tracking of one-color single-molecule data (using e.g. u-track (7)). Every object (monomer or oligomer) becomes a particle, and the only information stored are its position and intensity over time (as with experimental single-molecule data). When an association happens, two particles merge into one. When a dissociation happens, one particle splits into two. The time step for the output particle tracks is a user-specified parameter, generally chosen to match the experimental data time step.

Note that the oligomeric state of particles is not stored explicitly, as this information is not a direct output of single-molecule imaging experiments. However, for probe simulations, where the individual fluorophore intensity *I*_*ind*_ ∼ *N*(1,0), particle intensity is effectively equal to oligomeric state. For target simulations, on the other hand, the individual fluorophore *σ*_*ind*_ > 0, as would be the case for experimental single-molecule data. In this case, the oligomeric state of the particles is unknown and is subsequently estimated from the particle intensities and their merging and splitting history, as described in the next section.

### 4. Single-molecule data analysis to calculate intermediate statistics

#### Estimation of oligomeric state from particle intensities and merging and splitting history

The first step for calculating the intermediate statistics is to estimate the oligomeric state of each particle over time, i.e. it dynamic oligomerization history, using the available information, namely the particle intensities and their merging and splitting history. For this, particle tracks are divided into segments, where each segment starts via an appearance or a split, and ends via a disappearance or a merge. The dynamic oligomerization history is then determined as follows:

For probe data: Because the individual fluorophore intensity *I*_*ind*_ ∼ *N*(1,0) in probe simulations, the oligomeric state of a probe particle over time is given by its intensity over time.

For target data (whether experimental or simulated): The oligomeric state of each track segment is determined using constrained least squares minimization. Specifically, suppose a group of *N*_seg_ segments undergo merging and splitting events with each other (in the example shown in Fig S2A, N_seg_ = 5). Given a mean intensity *I*_*j*_ for each segment *j* (*j* = 1, 2, …, *N*_seg_), the oligomeric states *s*_*j*_ (*j* = 1, 2, …, *N*_seg_) of all segments in the group are determined by minimizing the least squares function *F*:

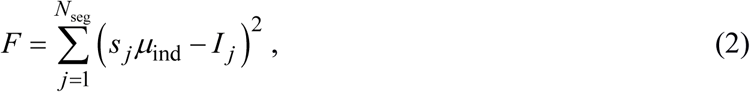

where *μ*_ind_ is the (known) mean intensity of an individual fluorophore. The minimization is subject to constraints from the merging and splitting history of the segments, imposing conservation of number of molecules. We will use the example shown in Fig S2A to illustrate the constraints: In that example, segments 1 and 2 merge to form segment 3, and then segment 3 splits into segments 4 and 5. This imposes the constraints that:

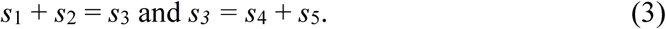

Thus, for the example shown in Fig S2A, *F* in Eq. 2 will be minimized subject to the constraints in Eq. 3. Given the integer nature of the unknown oligomeric states, the constrained least squares minimization is solved as a Mixed Integer Quadratic Programming problem (using the Package YALMIP (34) (https://yalmip.github.io/)). Of note, Eq. 2 is a simplification of the real objective function to be minimized, which should weight each term *j* by its oligomeric state, as the variance generally increases with number of fluorophores being summed up. In addition, fluorophore fluorescence can follow a log-normal distribution, rather than a normal distribution as assumed in Eq. 2 (35). Nevertheless, Eq. 2 is an acceptable simplification because the constraints play the major role in determining the oligomeric state.

This analysis yields the dynamic oligomerization history of the labeled molecules throughout the simulation/observation time.

#### Calculation of intermediate statistics

Within the framework of Indirect Inference, FISIK uses the labeled oligomer densities and the labeled molecule dissociation rates as intermediate statistics. These intermediate statistics were chosen because they are closely related to the unknown model parameters, namely the interaction parameters *p*_*a*_(*n*) and *k*_*off*_(*n*) (*n* = 2, 3, 4, …) and the population parameters *ρ* and *f*. Their choice was also validated a posteriori as they indeed enabled FISIK to estimate the unknown model parameters (as discussed in the Results section).

Using the labeled molecules’ dynamic oligomerization history (generated as described above), the intermediate statistics are calculated as follows for each simulation (or experimental time lapse in the case of experimental data):

##### (i) Labeled oligomer densities

Having the oligomeric state of every particle at every time point, the average density of monomers 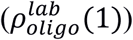, dimers 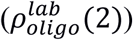, etc. is calculated. The superscript “lab” is to emphasize that these are the oligomeric states and their densities as revealed by the labeled subset of molecules.

##### (ii) Labeled molecule dissociation rates

These are derived by treating transitions between labeled oligomeric states as a Markov process. If there are labeled molecule oligomers up to oligomeric state 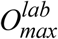, then the Markov process consists of 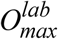 states (Fig S2B). For each oligomeric state *n* ≥ 2, we calculate from the dynamic oligomerization history its average lifetime *τ*(*n*) and its probability *p*_n→n-1_ to transition to state *n*-1 (i.e. a labeled molecule dissociates; Fig S2B). Inverting the Gillespie algorithm for the simulation of stochastic processes (36), the labeled molecule dissociation rate is then calculated as

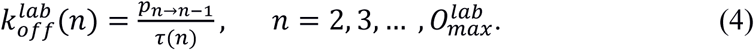

Note that, in order to minimize redundancy between intermediate statistics (21), the labeled molecule association rates are not used as intermediate statistics. This is because, under the assumption of steady state, they are simply calculated from the labeled molecule oligomer densities and labeled molecule dissociation rates:

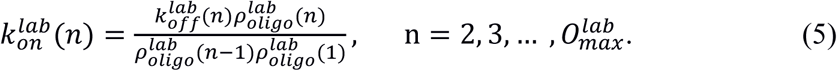

In other words, the labeled molecule densities and off rates are sufficient to capture all the available information about the molecular interactions of interest.

With this, the vector of intermediate statistics describing each simulation/experimental time lapse is constructed as

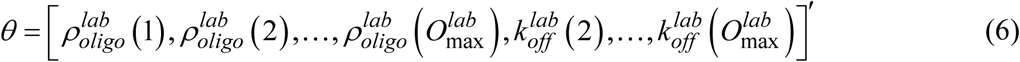

For a probe/target represented by multiple simulations (or multiple experimental time lapses), its vector of intermediate statistics *θ*_*prb*_/*θ*_*tar*_ and associated variance-covariance matrix *V*_*prb*_/*V*_*tar*_ are calculated as the mean and covariance, respectively, of the intermediate statistics vectors from its multiple simulations. The calculated *θ*_*prb*_ and *θ*_*tar*_ and associated *V*_*prb*_ and *V*_*tar*_ are then used in Eq. 1 to calculate the Mahalanobis distance to compare probes and targets.

## Results and Discussion

### The intermediate statistics accurately characterize the modeled molecular interactions

To validate our choice of intermediate statistics, we employed simulations where all molecules were labeled, in which case there are straightforward relationships between the input model parameters and the calculated intermediate statistics. We systematically varied the model parameters, i.e. *ρ, p*_*a*_, *k*_*off*_ and *D* and investigated how the intermediate statistics responded. Note that, for the sake of simplicity, in these and most of the following tests *p*_*a*_ refers to *p*_*a*_(2 ≤ *n* ≤ 5) (all equal) and *k*_*off*_ refers to *k*_*off*_(2 ≤ *n* ≤ 5) (also all equal). Unless explicitly stated otherwise, *p*_*a*_(*n* > 5) = 0, i.e. the maximum oligomer that could be formed was a pentamer. The simulation parameters employed for these tests are shown in Table S1, Row 1.

The calculated total molecule density (*ρ*^*lab*^) and dissociation rate 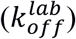 were equal to their corresponding input values, and they only varied when their input values were changed (Fig 2A-D, 1^st^ and 2^nd^ columns; 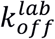 was slightly underestimated for input *k*_*off*_ = 2/s because of the employed output trajectory time step of 0.1 s). The distribution of oligomeric states 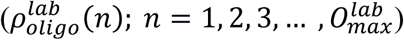 shifted toward higher values as the input *p*_*a*_ or *D* increased, or the input *k*_*off*_ decreased (Fig 2A-D, 1^st^ column). While not used as an intermediate statistic, we also investigated how the calculated association rate 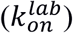 varied with model parameters, as this information is implicitly contained in 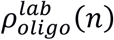 and 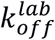. We found that 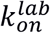 varied approximately linearly with the input *p*_*a*_ and *D*, and was largely independent of the input *ρ* and *k*_*off*_ (Fig 2A-D, 3^rd^ column), as it should be. These results validate our simulation and intermediate statistics calculation strategy, and support our choice of intermediate statistics to capture the relevant information about the system for model calibration.

Before proceeding to test FISIK, we also investigated how the intermediate statistics varied with the labeled fraction *f* (Fig 2E). This would aid us in interpreting our test results. As expected, reducing *f* shifted the distribution of labeled molecule oligomeric states toward lower values, completely eliminating in some cases the observation of higher oligomeric states, and reduced 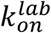. More interestingly, because less interaction events were visible for simulations with lower *f*, all oligomeric states, except for the highest one, appeared longer-lived than they truly were, leading to a reduced 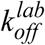 as well.

### FISIK can estimate interaction parameters from single-molecule trajectories with sub-stoichiometric labeling, provided a sufficient fraction/number of molecules is labeled

In order to achieve reasonably accurate particle tracking, most 2D single-molecule imaging experiments operate at a labeled molecule density roughly in the range 0.1-1 mol/μm^2^ (7, 9, 16). Therefore, as a first test of FISIK, we focused on simulations with relatively low molecule densities (*ρ* = 1-16 mol/μm^2^), in order to afford low-medium labeled fractions (*f* = 0.1-0.6). Taking the example of cell-surface receptors, such densities are realistic for various receptor types, such as CD36 (6), EpoR (37), LDLR (38) and various GPCRs (14, 17). Because the parameter inference problem is expected to get more difficult as the labeled fraction gets smaller – an expectation that is met in our tests as will be discussed shortly – these initial studies test FISIK under relatively favorable conditions.

For these studies we used the model parameters in Table S2, Row 1 (see also Fig 3) and simulation parameters in Table S1, Row 2. The values of *D* and *k*_*off*_ were motivated by single-molecule observations of a variety of cell surface receptors (6, 8, 14). The relatively small values of *p*_*a*_ were chosen to reflect that molecules must have the correct relative orientation in order to associate (39). Of note, the combination of *D* and *p*_*a*_ values indeed resulted in 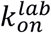 that was within the range of experimentally observed values (14, 17). The probe parameter landscape consisted of 3360 parameter combinations, each represented by 10 simulations. This probe landscape was tested against several targets, each also represented by 10 simulations. However, to mimic molecule density variability between cells, the target simulations differed from the probe simulations in that the density varied between the 10 simulations of each target, following a normal distribution with mean = the specified density and standard deviation = 0.1 × mean. Additionally, to mimic individual fluorophore intensity fluctuations and the fact that the oligomeric state is unknown in experimental single-molecule data, in the target simulations the individual fluorophore intensity fluctuated over time, following the distribution *I*_*ind*_ ∼ *N*(1,0.3) (in the probe simulations *I*_*ind*_ ∼ *N*(1,0)). The target parameters were chosen to span a wide range of values for the interaction parameters *p*_*a*_ and *k*_*off*_ and the population parameters *ρ* and *f*.

**Figure 3.**
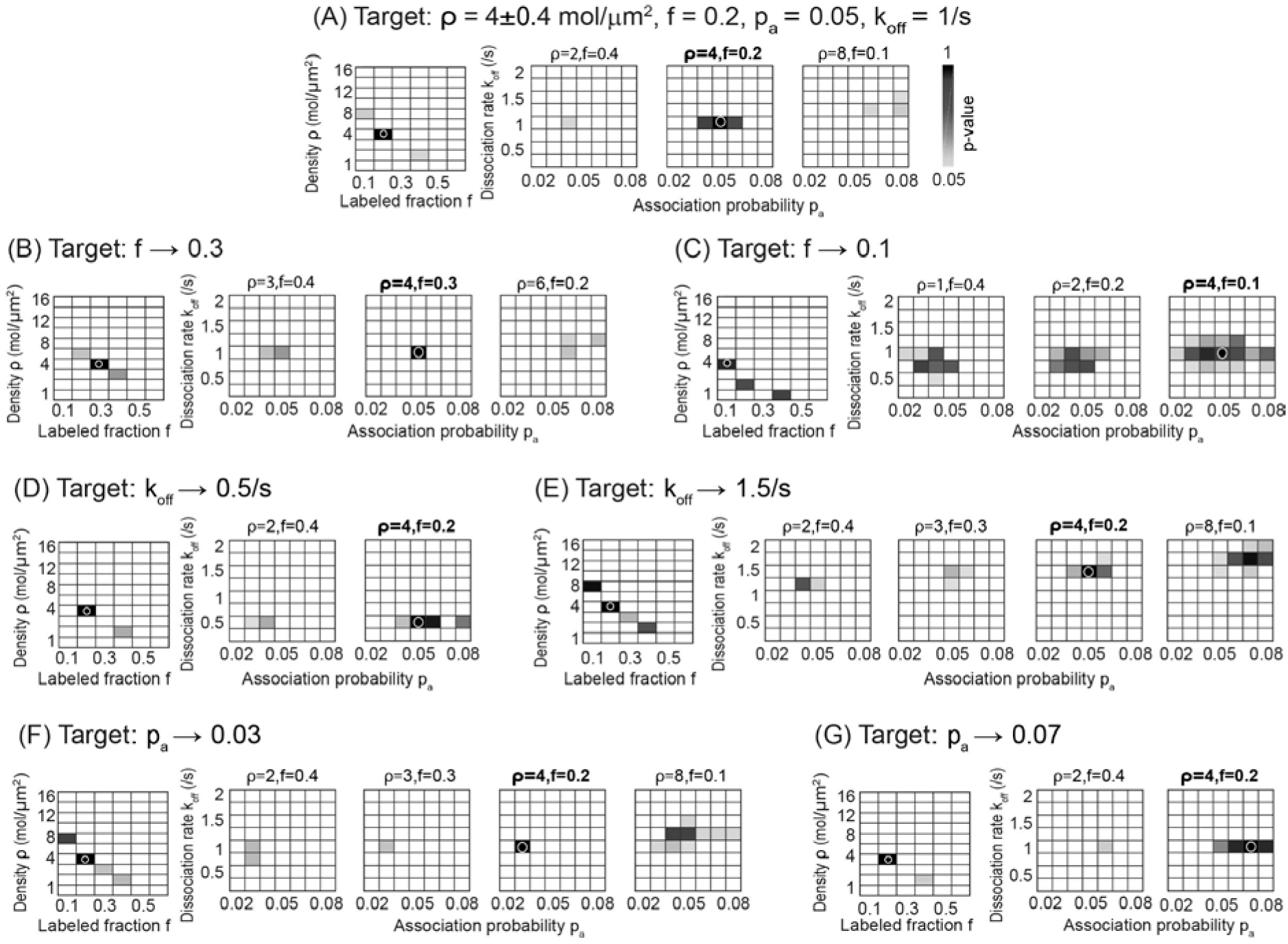
FISIK can estimate interaction parameters from single-molecule trajectories with sub-stoichiometric labeling, and can detect shifts in parameter values. (**A**) Four dimensional p-value landscape for comparing probe trajectories to the indicated target, as a function of the population parameters *ρ* and *f* and the interaction parameters *p*_*a*_ and *k*_*off*_ (total of 3360 probes). The p-value is shown as a heat map, going from white (p-value < 0.05) to black (p-value = 1). The *ρ* vs. *f* landscape is shown on the left, where each rectangle is a “summary view” of the *k*_*off*_ vs. *p*_*a*_ landscape corresponding to each *ρ-f* pair. Specifically, it shows the maximum p-value obtained for the underlying *k*_*off*_ vs. *p*_*a*_ landscape. The non-white rectangles in the *ρ* vs. *f* landscape are then “expanded” on the right, showing the corresponding full *k*_*off*_ vs. *p*_*a*_ landscapes. The specific *ρ-f* pairs being expanded are indicated above each *k*_*off*_ vs. *p*_*a*_ landscape, with the target *ρ-f* pair in bold. The target parameter values are also indicated by a white open circle in the *ρ* vs. *f* landscape on the left and a white open circle in the *k*_*off*_ vs. *p*_*a*_ landscape on the right. (**B-G**) Same as (A), but with target *f* increased (**B**) or decreased (**C**), target *k*_*off*_ decreased (**D**) or increased (**E**), or target *p*_*a*_ decreased (**F**) or increased (**G**) to the indicated values. The correct probe parameters (= target parameters) had the following p-values: 0.9967 (rank 1/7 possible solutions) in (A), 0.9915 (rank 1/6 possible solutions) in (B), 0.8120 (rank 4/30 possible solutions) in (C), 0.9998 (rank 1/6 possible solutions) in (D), 0.9963 (rank 1/17 possible solutions) in (E), 0.9995 (rank 1/14 possible solutions) in (F), and 0.9871 (rank 1/5 possible solutions) in (G).

In the following we discuss our major findings from these tests, using representative targets for demonstration. Taking the target *ρ* = 4 ± 0.4 mol/µm^2^, *f* = 0.2, *p*_*a*_ = 0.05 and *k*_*off*_ = 1/s as our reference point, we will first explain how the parameter inference results are displayed (Fig 3A). As explained in Methods, Section 1, the difference between a probe and a target is calculated as the Mahalanobis distance between their intermediate statistics (Eq. 1), which is converted to a p-value. Thus the results are shown as p-value landscapes. However, because we are searching in a 4-dimensional parameter space, the p-value landscapes are shown in two steps. In the first step, the *ρ* vs. *f* landscape is shown on the left, where each rectangle is a “summary view” of the *k*_*off*_ vs. *p*_*a*_ landscape corresponding to each *ρ*-*f* pair (specifically, it shows the maximum p-value obtained for the underlying *k*_*off*_ vs. *p*_*a*_ landscape). White rectangles indicate *ρ*-*f* pairs containing no matches with the target (for any tested *k*_*off*_-*p*_*a*_ combination). Non-white rectangles indicate *ρ-f* pairs containing some matches with the target (for some *k*_*off*_*-p*_*a*_ pairs, as explained next). Darker rectangles indicate matches with higher p-value. In the second step, the non-white rectangles in the *ρ* vs. *f* landscape are “expanded” on the right, to show the full *k*_*off*_ vs. *p*_*a*_ landscapes corresponding to those *ρ-f* pairs. Again, white rectangles indicate *k*_*off*_*-p*_*a*_ pairs (for each *ρ-f* pair) that do not match the target (p-value < 0.05), non-white rectangles indicate *k*_*off*_*-p*_*a*_ pairs (for each *ρ-f* pair) that are considered to match the target (p-value ≥ 0.05), and darker rectangles indicate higher p-values.

For the reference target (Fig 3A), FISIK was able to identify the correct parameters (indicated by white circles; p-value = 0.9967), within a small range of solutions. This range of solutions was in part due to the expected coupling between the population parameters *ρ* and *f*, such that the estimated *ρ* × *f* ≈ the average density of labeled molecules in the target (= 0.8 labeled mol/μm^2^ for this target). Note, however, that this coupling did not extend indefinitely. For example, comparing the probe landscape to a target with *ρ* = 100 ± 10 mol/µm^2^, *f* = 0.03, *p*_*a*_ = 0.05 and *k*_*off*_ = 1/s (described in detail in the next section) did not yield any matches between them, even though 100 × 0.03 = 3 labeled mol/µm^2^, a labeled molecule density that was achievable by multiple *ρ-f* combinations in the probe landscape. We suspect that the finite range of *ρ-f* combinations in the solution is due to constraints imposed by estimating these two population parameters in the context of the interaction parameters *k*_*off*_ and *p*_*a*_. Therefore, even though FISIK yields a range of estimated *ρ* and *f* values, the range is finite and contained within one order of magnitude at most.

In spite of the observed *ρ*-*f* coupling, which reduced the determinability of *ρ* and *f*, the estimated *k*_*off*_ and *p*_*a*_, i.e. the interaction parameters of interest, were relatively consistent for the different *ρ-f* combinations. Both exhibited a narrow range, with high p-value solutions at *k*_*off*_ = 1/s and *p*_*a*_ = 0.04-0.06 (target *k*_*off*_ = 1/s and *p*_*a*_ = 0.05). The low p-value solutions at *ρ* = 8 and *f* = 0.1 were slightly shifted up for both *k*_*off*_ (1.25-1.5/s) and *p*_*a*_ (0.06-0.08), most likely to compensate for the lower estimated 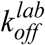 and 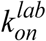 for lower *f* (Fig 2E). Testing FISIK against targets of different population parameters *f* and *ρ* in the neighborhood of the reference parameters indicated that targets with higher *f* or *ρ* led to narrower solution ranges, while targets with lower *f* or *ρ* led to wider solution ranges, reducing the determinability of some parameters, especially *p*_*a*_ (Fig 3B, C and Fig S3).

Next we tested the sensitivity of FISIK to detect shifts in interaction rates. Varying target *k*_*off*_ from 1/s to 0.5/s or to 1.5/s led to clear shifts in the solution range, from 1-1.5/s for target *k*_*off*_ = 1/s to 0.5/s for target *k*_*off*_ = 0.5/s and to 1.25-2/s for target *k*_*off*_ = 1.5/s (Fig 3A, D, E). Varying *p*_*a*_ from 0.05 to 0.03 or to 0.07 shifted the high p-value solution range from 0.04-0.06 for target *p*_*a*_ = 0.05, to 0.03-0.05 for target *p*_*a*_ = 0.03, and to 0.06-0.08 for target *p*_*a*_ = 0.07 (Fig 3A, F, G).

All in all, these tests demonstrate that FISIK is able to estimate molecular interaction parameters from single-molecule data, as long as there is a sufficient fraction/number of labeled molecules. Because of the slight dependence of the estimated interaction parameters on the estimated population parameters, combining FISIK with prior knowledge about the molecule density, which is experimentally achievable via methods such as flow cytometry (40, 41), is expected to aid FISIK with this task.

### The fraction of labeled molecules in single-molecule trajectory data is a critical factor for FISIK’s ability to estimate interaction parameters

While many transmembrane proteins exist at a surface density in the 1-10 mol/μm^2^ range, others are much more highly expressed. Examples of receptors with high surface densities are CD8 (42) and receptors of the ErbB family (38, 43). In such cases, the labeled fraction *f* will be below 0.1 to maintain single-molecule trackability. Therefore, as a next test of FISIK, we simulated targets and probes with *ρ* ≈ 100 mol/μm^2^ and *f* close to 0.1 and below. These tests not only investigated the performance of FISIK when *f* is very low, but also allowed us to determine more conclusively which property is more important: the labeled fraction *f* or the labeled molecule density = *ρ* × *f* (the tests above showed some interdependence between the minimum necessary *f* and *ρ*; see Fig 3A, C and Fig S3C).

For these studies we used the model parameters in Table S2, Row 2 and the simulation parameters in Table S1, Row 3. The probe parameter landscape consisted of 2352 parameter combinations, where each parameter combination was represented by 30 simulations (here the simulation area was smaller for the sake of simulation efficiency, and thus we compensated for the smaller area, i.e. smaller number of events, by combining 30 simulations instead of 10). The target parameters were *ρ* = 100 ± 10 mol/µm^2^, *p*_*a*_ = 0.05, *k*_*off*_ = 1/s and multiple values for *f* to investigate its effect on interaction parameter determinability (also 30 simulations per target).

First we investigated the solution range when the target *f* = 0.03 (Fig 4A), because a target with *ρ* = 100 ± 10 mol/µm^2^ and *f* = 0.03 has the same labeled molecule density as a target with *ρ* = 10 ± 1 mol/µm^2^ and *f* = 0.3, a target for which FISIK yielded an almost perfect solution (Fig S3D). However, in contrast to the *ρ* = 10 ± 1 mol/μm^2^ and *f* = 0.3 case, the *ρ* = 100 ± 10 mol/μm^2^ and *f* = 0.03 case had a wide range of solutions, with *p*_*a*_ practically undeterminable. This test clearly demonstrates that the labeled fraction *f*, and not the labeled density per se, is the most critical parameter influencing the performance of FISIK.

**Figure 4.**
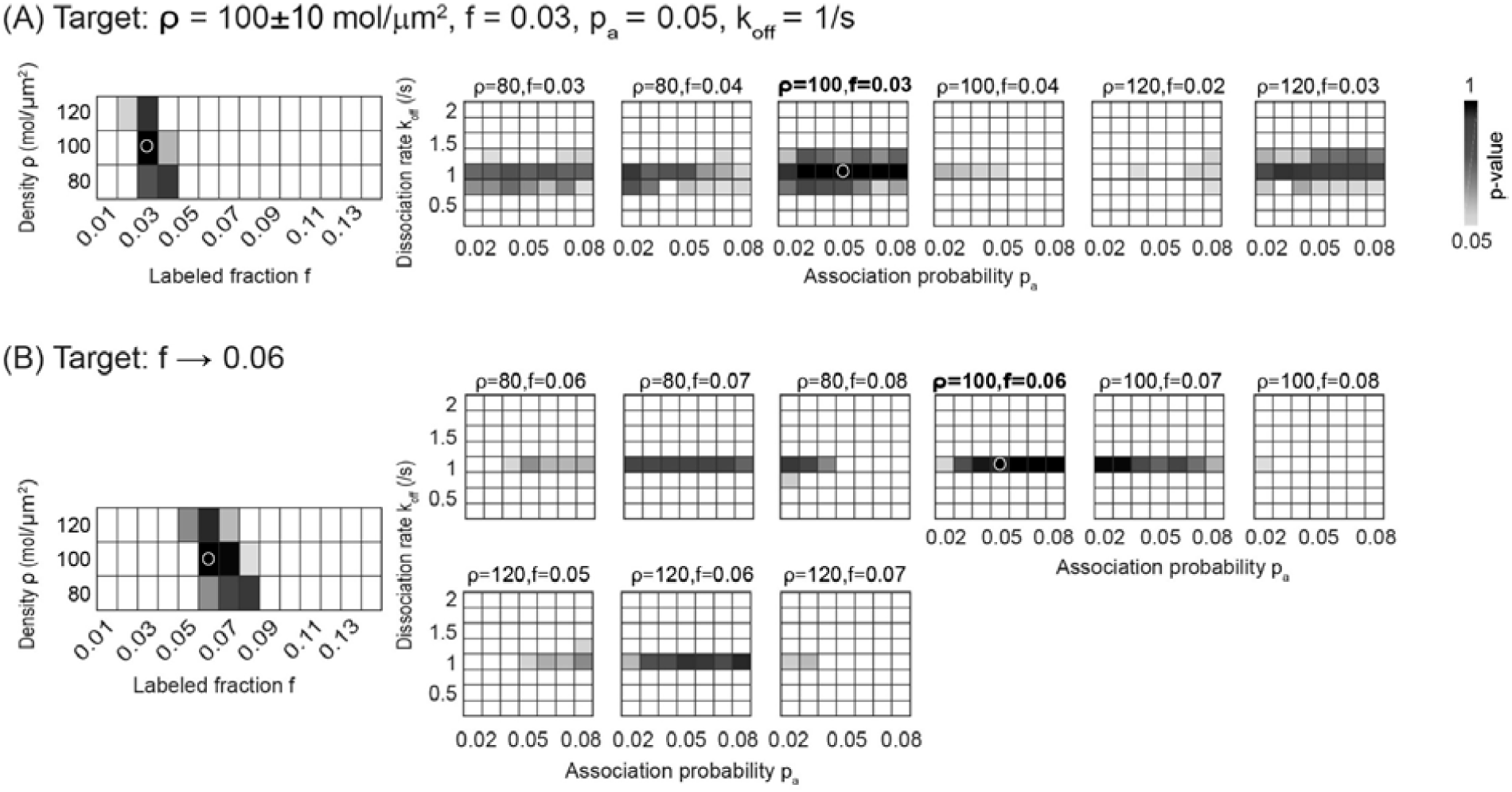
The dissociation rate *k*_*off*_ can be estimated accurately with a smaller fraction of labeled molecules than the association probability *p*_*a*_. (**A**) Four dimensional p-value landscape for comparing probe trajectories to the indicated target, as a function of the population parameters *ρ* and *f* and the interaction parameters *p*_*a*_ and *k*_*off*_ (total of 2352 probes). Detailed description as in Fig 3A. (**B**) Same as (A), but with target *f* increased to 0.6. The correct probe parameters (= target parameters) had the following p-values: 0. 9880 (rank 2/87 possible solutions) in (A), and 0.91 (rank 6/45 possible solutions) in (B).

To determine the minimum labeled fraction needed to accurately estimate the interaction parameters, next we increased the target *f* from 0.03 to 0.06 (Fig. 4B), 0.09 and 0.12 (data not shown). As *f* increased, the solution range decreased. By *f* = 0.06, *k*_*off*_ was estimated accurately within a narrow range (all solutions were at 1/s; Fig 4B). The range of *p*_*a*_ also gradually decreased as *f* increased, although it was still quite wide even at *f* = 0.12 (data not shown).

Putting together the results of the tests here and in the previous section, we conclude that the labeled fraction of molecules is a critical parameter for FISIK’s performance. The number/density of labeled molecules is important (e.g. Fig 3A vs. Fig S3C), but the labeled fraction of molecules is much more critical. When the labeled fraction is too small, a too large fraction of multi-molecular events in the system, such as association, becomes invisible. Fundamental information is lost about these multi-molecular processes, and the information seems to be irrecoverable. As a result, determining *p*_*a*_, which characterizes a mutli-molecular process (association), requires labeled fractions of at least ∼0.2 (Fig 3A). In contrast, determining *k*_*off*_ is possible with relatively low labeled fractions, down to ∼0.06 (Fig 4B), as dissociation is a uni-molecular process.

### Static oligomer distribution data can complement low-labeled-fraction single-molecule trajectory data and increase FISIK’s power

Our results thus far pose a dilemma. The inference of interaction parameter values, especially *p*_*a*_, improves as the labeled fraction of molecules increases. Yet increasing the labeled fraction often increases particle tracking errors (7, 16), thus reducing the data quality for model calibration and parameter inference (the effect of tracking errors will be investigated in more detail later). One approach to address this challenge is to employ multi-color single-molecule imaging (e.g. 2- or 3- color, or even more with hyperspectral imaging (4)), where the labeled fraction in each channel affords good trackability, while the sum of all channels gives a high enough total labeled fraction. This approach is conceptually similar to the above tests, and thus will not be explored further here. Another approach to address this challenge is to combine the dynamic but sparse information from single-molecule tracking with dense but static information from approaches such as Number and Brightness (N&B) analysis (44, 45), Spatial Intensity Distribution Analysis (SpIDA) (46), and Single-Molecule Localization Microscopy (SMLM) (47, 48). These approaches yield a dense snapshot of the oligomeric state distribution of the majority of molecules of interest, thus potentially complementing low-labeled-fraction single-molecule tracking data and improving the ability of FISIK to infer interaction parameters.

As a first step, we investigated the performance of FISIK when using only static oligomer distribution data. For these tests we used the model parameters in Table S2, Row 3 and the simulation parameters in Table S1, Row 4. In terms of model parameters (Table S2), we replaced the labeled fraction f with the observed fraction *ϕ*. The observed fraction accounts for the fact that, even when all molecules are labeled, not all molecules are observed, due to incomplete fluorophore maturation (17), or incomplete photoconversion and localization in the case of SMLM (49, 50). In the targets we used *ϕ* = 0.85, and made *ϕ* an unknown parameter to estimate; in the probes *ϕ* was varied in the range 0.75-0.95 (17, 49, 50). The output of the static snapshot simulations was the particle intensities and positions at the end of the simulation, equivalent to the output of the full trajectory simulations, but for one-time point (Methods, Section 3). As there is no merging and splitting history in the static snapshot data, the oligomeric state of each particle in this case was determined solely from the particle intensities, i.e. by minimizing the objective function in Eq. 2 but without any constraints (Methods, Section 4). Additionally, the only intermediate statistics to be used for target and probe comparison in the case of static distribution data were the observed oligomer densities: 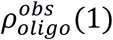 (monomers), 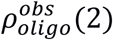 (dimers), etc.

These tests demonstrated that implementing FISIK with static oligomer distribution data instead of dynamic single-molecule trajectory data yielded an accurate estimate of the ratio of *k*_*off*_ to *p*_*a*_ (Fig 5A). Note that *p*_*a*_ is linearly related to the association rate *k*_*on*_ (Fig 2C); as a result, *k*_*off*_ /*p*_*a*_ ∝ *k*_*off*_ /*k*_*on*_ = *K*_*d*_, the dissociation constant. In other words, applying FISIK with dense, static oligomer distribution data extracts from these data the maximum information possible, namely the dissociation constant(s) describing molecular interactions. The accuracy of the estimated *K*_*d*_’s is expected to depend on the accuracy of the experimental and analytical procedures employed to calculate the distribution of oligomeric states. For example, in the case of SMLM, fluorophore blinking and multiple appearances must be accounted for properly to avoid over-counting that results in exaggerated apparent oligomerization (47).

**Figure 5.**
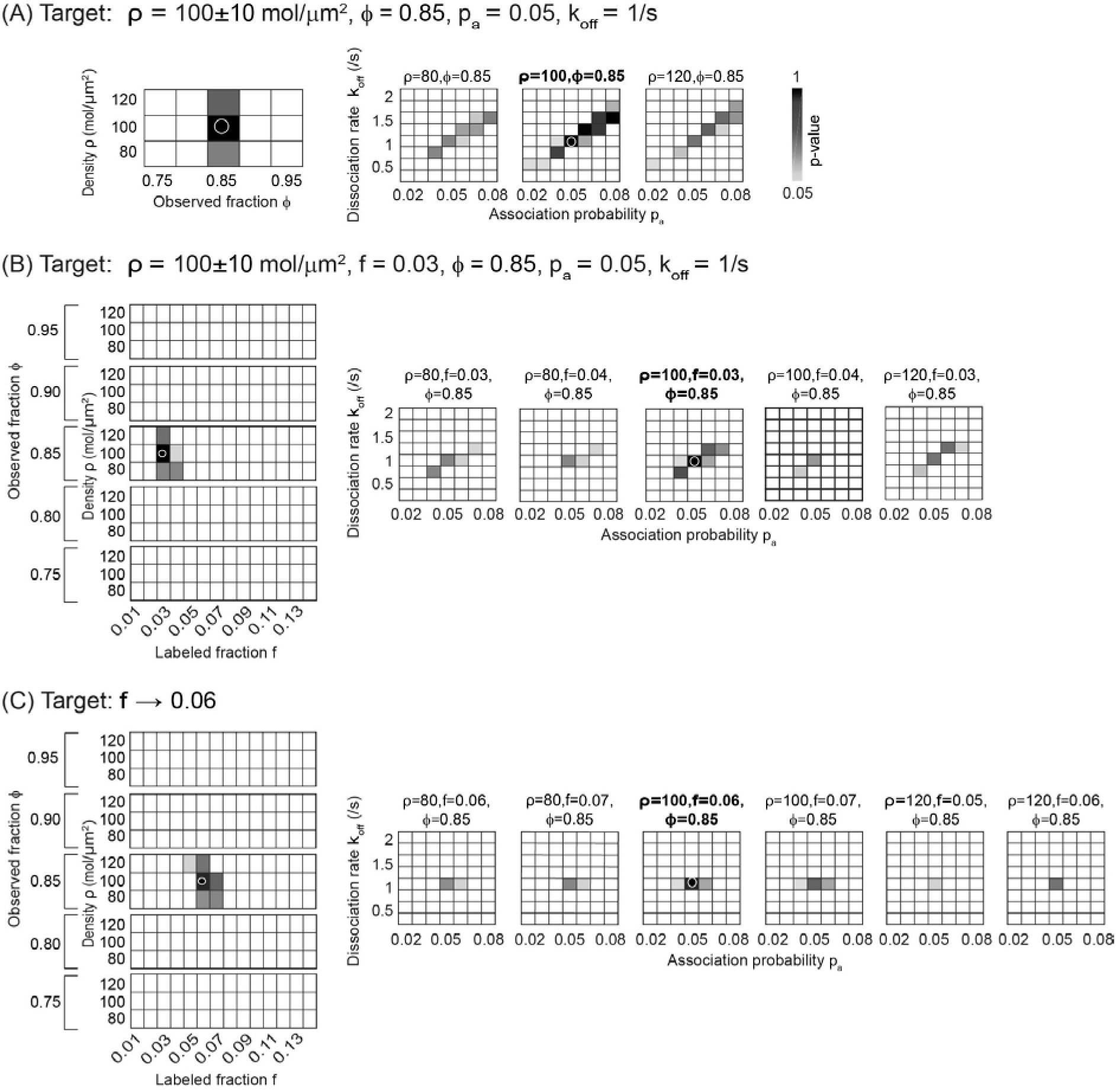
Dense but static oligomer distribution data increase FISIK’s power in the case of sparse single-molecule trajectory data. (**A**) Four-dimensional p-value landscape for comparing probe oligomer distribution data to the indicated target oligomer distribution data, as a function of the population parameters ρ and *ϕ* and the interaction parameters *p*_*a*_ and *k*_*off*_ (total of 840 probes). Detailed description as in Fig 3A, except that the labeled fraction *f* is replaced by the observed fraction *ϕ* (see text for details). (**B**) Five-dimensional p-value landscape for comparing combined probe data (trajectories + distribution data) to the indicated target (also trajectories + distribution data), as a function of the population parameters *ρ, f* and *ϕ* and the interaction parameters *p*_*a*_ and *k*_*off*_ (total of 11760 probes). Detailed description as in Fig 3A, except that the parameter space includes both *f* and *ϕ*. (**C**) Same as (B), but with target *f* increased to 0.6. The correct probe parameters (= the target parameters) had the following p-values: 0.9995 (rank 1/28 possible solutions) in (A), 0.9980 (rank 1/23 possible solutions) in (B), 0.91 (rank 1/12 possible solutions) in (C).

Our tests thus far indicate that FISIK with low-labeled-fraction single-molecule trajectory data yields accurate estimates of *k*_*off*_, while FISIK with static oligomer distribution data yields accurate estimates of *k*_*off*_/*p*_*a*_. Therefore, we surmised that combining the two data types should yield an accurate estimate of both *k*_*off*_ and *p*_*a*_, along with estimating *ρ, f* and *ϕ* (i.e. 5 parameters in total). We implemented the combination of data types at the level of FISIK’s solution space. Specifically, we defined the combined p-value for probe-target comparison as:

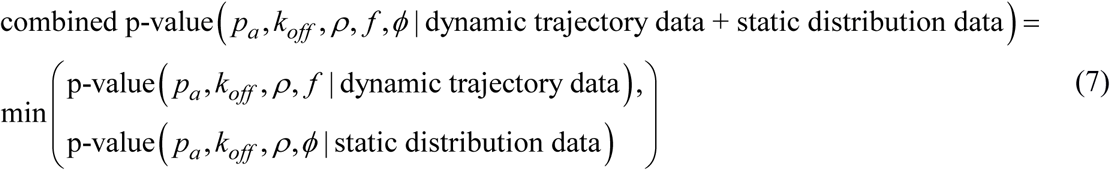

where the p-values are calculated from the Mahalanobis distance as done previously (Eq. 1). With this, a combination of *p*_*a*_, *k*_*off*_ and *ρ* values will only be accepted as a possible solution if it is a possible solution in both the dynamic comparison and the static comparison. This strategy indeed enabled the accurate estimation of *p*_*a*_ and *k*_*off*_ with a single-molecule labeled fraction as low as 0.06 (Fig 5B, C; for *f* = 0.06, estimated *p*_*a*_ range 0.04-0.06 and *k*_*off*_ = 1/s for target *p*_*a*_ = 0.05 and *k*_*off*_ = 1/s). These tests demonstrate that combining these two types of complementary data is indeed a viable approach to enhance the accuracy of FISIK, without the need to increase the labeled fraction in single-molecule trajectory data.

### FISIK can estimate molecular interaction parameters in the presence of limited mismatch between model and system

In our tests of FISIK thus far, there was no mismatch between model (i.e. probe) and the system it mimics (i.e. target). However, when applying FISIK to experimental data, it is highly likely that the employed model is an approximation of reality. Therefore, we investigated the performance of FISIK when there is a model mismatch between probe and target. Specifically, we investigated the effect of a mismatch in the maximum achievable oligomeric state (*O*_*max*_, such that *p*_*a*_(*n*>*O*_*max*_) = 0)), and a mismatch in the diffusion coefficient D, as both of these are parameters that we have thus far assumed known instead of determining their values through model fitting.

To test the effect of mismatches in *O*_*max*_, we simulated targets with different *O*_*max*_, while retaining the probe *O*_*max*_ at 5 (Fig 6A). The employed target parameters were otherwise those of our reference target (*ρ* = 4 ± 0.4 mol/µm^2^, *f* = 0.2, *p*_*a*_ (2≤n≤ *O*_*max*_) = 0.05 and *k*_*off*_ = 1/s; Fig 3A). High p-value solutions started to appear at target *O*_*max*_ = 3, where the interaction parameters were estimated with reasonable accuracy (high p-value solutions for *p*_*a*_ = 0.04-0.07 and *k*_*off*_ = 1.25-1.5/s), although at the wrong *ρ*-*f* combination (8×0.1 instead of 4×0.2). Increasing *O*_*max*_ to 4, the interaction parameters were estimated accurately and at the correct *ρ*-*f* combination, although high p-value solutions were still present at the 8×0.1 *ρ*-*f* combination. For *O*_*max*_ > 5, the solution range was very similar to that for target *O*_*max*_ = 5 (Fig 6A vs. Fig 3A). These results suggest that FISIK’s performance is relatively robust against mismatches in *O*_*max*_; if it is able to estimate the interaction parameters, they are estimated reasonably accurately. If the mismatch in *O*_*max*_ between target and probe is too large, it appears that FISIK does not yield any solution, instead of an erroneous one. This robustness however implies that FISIK as applied to single-molecule tracking data alone is not able to estimate *O*_*max*_. Other techniques, such as static distribution data, would be better suited to estimate *O*_*max*_, which should then be input into the model.

**Figure 6.**
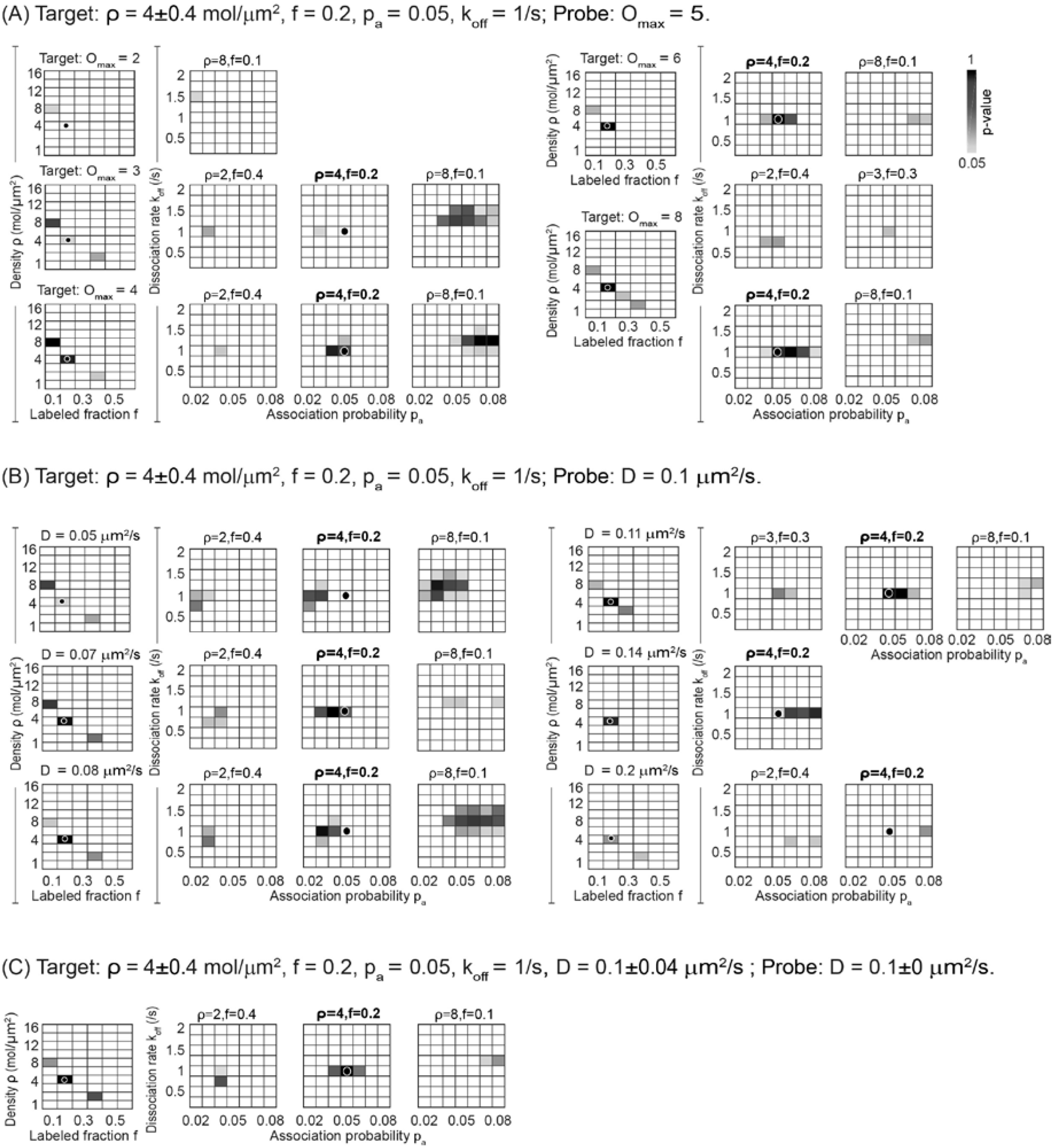
FISIK can estimate molecular interaction parameters in the presence of limited model mismatch between probe and target. Four dimensional p-value landscape for comparing probe trajectories to the indicated targets, as a function of the population parameters *ρ* and *f* and the interaction parameters *p*_*a*_ and *k*_*off*_ (total of 3360 probes). Targets are subject to mismatch in the maximum achievable oligomeric state *O*_*max*_ (**A**), and in the diffusion coefficient *D* (**B-C**). In (B), all molecules in target have the indicated *D* (which is different from the probe *D*). In (C), *D* for target molecules follows the indicated normal distribution. Detailed description as in Fig 3A. In (A), the correct probe parameters (= target parameters) were outside of the *O*_*max*_ = 2 and *O*_*max*_ = 3, and had the following p-values for the other tests: 0.6418 (rank 5/13 possible solutions) for *O*_*max*_ = 4, 0.9892 (rank 1/5 possible solutions) for *O*_*max*_ = 6, and 0.8984 (rank 2/10 possible solutions) for *O*_*max*_ = 8. In (B), the correct probe parameters were outside of the solution range for *D* = 0.05, 0.07, and 0.2 µm^2^/s, and had the following p-values for the other tests: 0.5165 (rank 3/10 possible solutions, for *D* = 0.08 µm^2^/s), 0.9738 (rank 2/8 possible solutions, for *D* = 0.11 µm^2^/s), 0.1061 (rank 4/4 possible solutions, for *D* = 0.14 µm^2^/s) in (B). In (C), the correct probe parameters had a p-value = 0.9991 (rank 1/7 possible solutions).

To test the effect of mismatches in *D*, we simulated targets with different *D*, while retaining the probe *D* at 0.1 µm^2^/s. Again, the employed target parameters were otherwise those of our reference target (Fig 3A). We found that lower/higher *D* in target compared to probe were compensated for by lower/higher *p*_*a*_ in the solution (Fig 6B). This is most likely because of the solution range for approximately linear dependence of 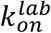 on *p*_*a*_ and *D* (Fig 2C and D, 3^rd^ column). Mismatches in *D* of ∼20% were tolerated (e.g. target D = 0.08 or 0.11 µm^2^/s), but beyond that the estimated *p*_*a*_ range no longer contained the true target *p*_*a*_, at least at the correct *ρ*-*f* combination. The tested mismatches in *D* had very little effect on estimating *k*_*off*_, on the other hand. In another test, we allowed *D* to vary between molecules in the target, following a normal distribution with mean = probe *D* (0.1 µm^2^/s), while retaining that all molecules in the probe had the same *D*. Even up to a standard deviation = 40% of the mean *D* (for larger standard deviations the distribution shifts substantially from a normal distribution, as *D* cannot be negative), the solution range remained very similar to that with no variation in *D* between molecules (Fig 6C vs. Fig 3A).

These tests indicate that FISIK is relatively sensitive to mismatches in the mean *D* between model and system, where mismatches in *D* are compensated for with the estimated *p*_*a*_, but it is robust against heterogeneity between molecules around their mean *D*. Therefore, especially in light of the coupling between *p*_*a*_ and *D* that these tests revealed, it is important to obtain good estimates of the mean *D*, potentially as a function of oligomeric state, via thorough analysis of the single-molecule trajectories, to use in the model. Yet the heterogeneity around the mean *D* does not have to be explicitly modeled.

### FISIK can estimate molecular interaction parameters from single-molecule trajectories subject to resolution limitations and tracking errors

While single-molecule imaging can reveal colocalization of molecules, suggestive of interactions, on its own it cannot distinguish between molecules interacting with vs. passing by each other, largely because of the resolution limit. Note that for the purposes of this section we are using the term interactions in a general sense, thus including direct interactions, indirect interactions and clustering within a common nanodomain. Distinguishing between these different types of interactions depends on the known biology and further experimentation of the system under study. Here we are concerned with distinguishing between them and coincidental colocalization due to molecules passing by each other. One-color single-molecule imaging suffers the most from the limited resolution of light microscopy. Multi-color imaging suffers less due to the ability to use e.g. co-movement in addition to colocalization to assess specific interactions (8). FRET-based experiments suffer the least from the resolution limit, due to the requirement of very close proximity for FRET. Nevertheless, fundamentally all light microscopy approaches face the challenge of distinguishing between proximity due to interactions vs. due to passing-by events.

Here we tested FISIK’s performance when applied to single-molecule trajectories obtained from one-color imaging data, as the most challenging scenario. We and others have previously developed particle tracking algorithms to capture merging and splitting events between imaged molecules in one-color time-lapses (7, 51, 52), reflecting molecular interactions (in the general sense defined above) (6, 13). For these tests, we used low density and low-medium labeled fraction simulations (Table S2, Row 1), as FISIK had an overall good performance under these conditions in the absence of particle tracking errors (Fig 3). We generated synthetic single-molecule image series that mimicked experimental time-lapses from simulated target trajectories (see Table S3 for image generation parameters). Then we detected and tracked the molecules in the synthetic image series using u-track (7), as we would detect and track actual experimental data (see Tables S4 and S5 for detection and tracking parameters). Finally, we used the tracking output, after a minor “clean up” (Note S1), as the target data for model calibration within FISIK. Note that the probe trajectories were used directly as output by the Monte Carlo simulations.

With the above strategy, we tested FISIK by generating images with an average signal-to-noise ratio (SNR) of 20 (high) or 7 (medium). High SNR conditions are not very common in single-molecule data, but they are achievable with quantum dot labeling (6, 8), for example. Medium SNR conditions are achievable with many organic dyes imaged via total internal reflection fluorescence microscopy (TIRFM), a common combination for single-molecule imaging. We started with a target with *ρ* = 2 ± 0.2 mol/μm^2^, *f* = 0.2, *p*_*a*_ = 0.05 and *k*_*off*_ = 1/s (as in Fig S3C). We chose this *ρ* and *f* combination because a labeled-molecule density of 0.4 mol/μm^2^ afforded (visually) acceptable detection and tracking performance (Videos S1-S4), and our previous tests on error-free data showed that *f* should be at least 0.2 for FISIK to estimate *p*_*a*_ well.

For both SNRs, FISIK estimated *k*_*off*_ well, within a narrow range, albeit slightly shifted toward smaller *k*_*off*_ values (estimated *k*_*off*_ in the range 0.5-1/s, with most high p-value solutions at 0.5/s and 0.75/s; Fig 7A). We surmised that this was due to the tracking software constraint that merges and splits were only allowed when there was no possibility of gap closing (Table S5). This constraint helped reduce the assignment of merge-to-split events to molecules merely passing by each other, but at the same time it might eliminate short-lived specific interactions between molecules. To investigate this issue further, we tested FISIK against targets with lower and higher *k*_*off*_ values. Indeed, the downward shift was more pronounced for higher target *k*_*off*_ values (Fig 7B, C). These observations imply that FISIK can estimate *k*_*off*_ accurately from single-molecule imaging data subject to resolution limitations, as long as the dissociation time scale is considerably longer than the time scale of molecules diffusing past each other, in order to distinguish between the two events.

**Figure 7.**
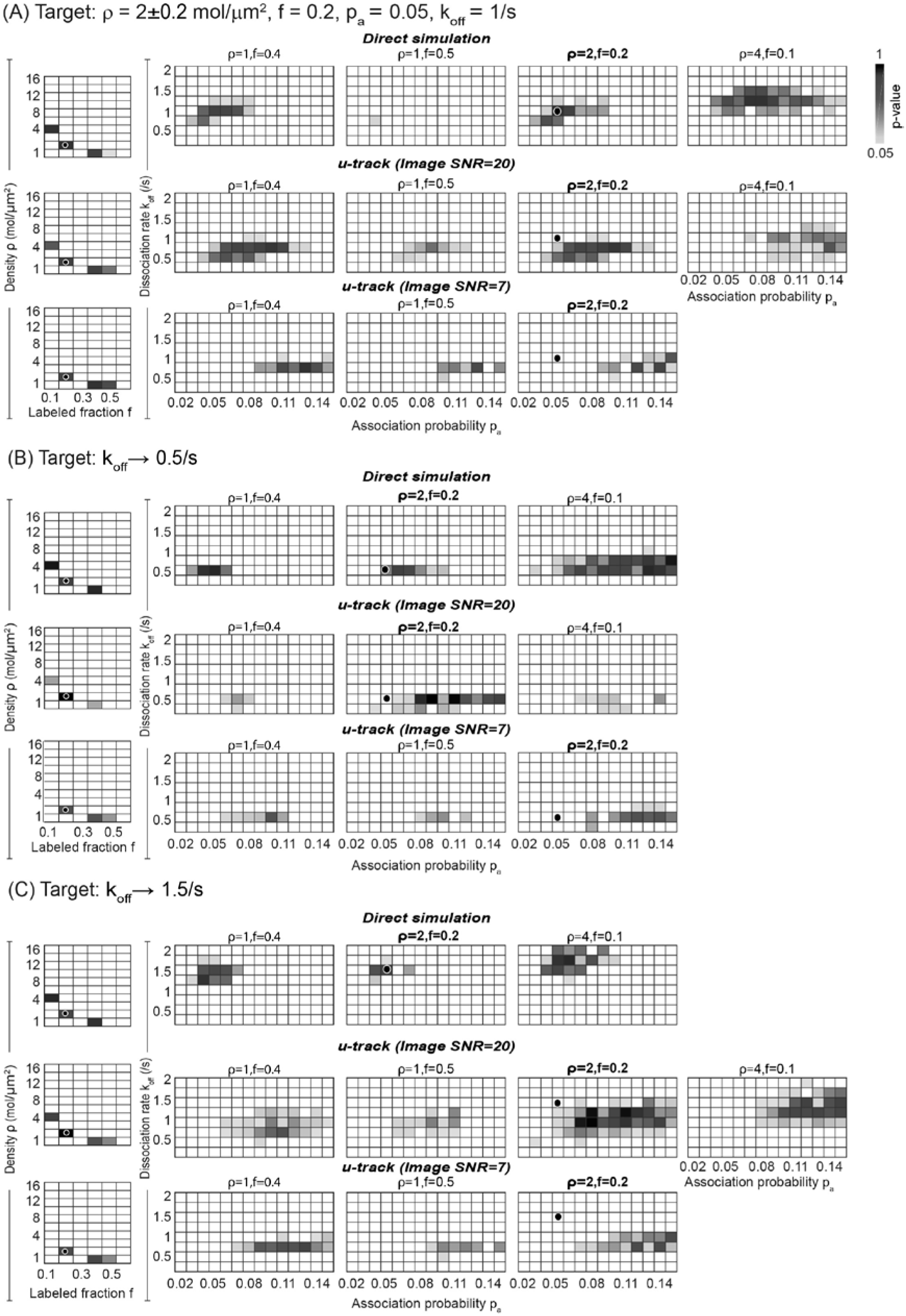
FISIK’s estimation of interaction parameters from single-molecule trajectories is influenced by resolution limitation and image SNR. (**A**) Four dimensional p-value landscape for comparing probe trajectories to the indicated target, as a function of the population parameters *ρ* and *f* and the interaction parameters *p*_*a*_ and *k*_*off*_ (total of 3584 probes; see Table). Detailed description as in Fig 3A. Top row (titled “Direct simulation”) shows landscape for comparing to target trajectories directly as output by the Monte Carlo simulations (as done for all figures up to Fig 6). Middle and bottom rows (titled “u-track (image SNR=20)” and “u-track (image SNR=7),” respectively) show landscape for comparing to target trajectories as output by u-track analysis of images with indicated SNR (see text for image generation and tracking details). (**B, C**) Same as (A), but with *k*_*off*_ decreased (**B**) or increased (**C**) to the indicated values. For direct simulation comparisons, the correct probe parameters (= target parameters) had the following p-values: 0.8917 (rank 2/72 possible solutions) in (A), 0.4138 (rank 14/49 possible solutions) in (B), and 0.7717 (rank 14/47 possible solutions) in (C). For u-track comparisons, the correct parameters had a p-value < 0.05 and were outside of the solution range.

Estimating *p*_*a*_ was more challenging than estimating *k*_*off*_, and depended more strongly on the image SNR. Specifically, the range of solutions for *p*_*a*_ was shifted toward higher values, and the shift increased as the SNR decreased. For SNR = 20, the target *p*_*a*_, was often just at the periphery of the solution range (Fig 7A-C, middle rows). For SNR = 7, the solution was shifted further away from the target *p*_*a*_ of 0.05, and was generally in the range 0.09-0.15 (Fig 7A-C, bottom rows). Close inspection of the detection and tracking steps of u-track suggested that FISIK performed worse with lower SNR because of detection errors (specifically related to the detection of very nearby particles) that propagated into tracking errors, as the tracking step had to balance the potentially conflicting events of gap closing, merging and splitting. Nevertheless, the solution range for *p*_*a*_ was finite and within 2-4-fold of the target *p*_*a*_, at least for average SNRs down to 7.

The requirement for relatively high SNR raised the concern of the effect of photobleaching on FISIK’s performance. To address this issue, we simulated targets where the fraction of labeled molecules decreased over time to mimic photobleaching. Starting with our reference target (Fig 3A), we simulated a range of photobleaching rates, inspired by TIRF-based single-molecule imaging using organic dyes (Fig S4A). In contrast to the targets, the probes did not include any photobleaching. As expected, for low photobleaching rates that retained ∼90% of the labeled molecules by the end of a 20 s “time-lapse” (0.005 and 0.007/s in Fig S4A), FISIK’s performance was very similar to that in the absence of photobleaching (Fig S4B, left, vs. Fig 3A). As the photobleaching rate increased further, such that less than 50% of labeled molecules were retained by the end of a 20 s “time-lapse” (0.03 and 0.05/s in Fig S4A), the solution range shifted toward lower *ρ* × *f* combinations, presumably to compensate for photobleaching (Fig S4B, bottom row). The interaction parameters remained in the vicinity of their target values, although *p*_*a*_ became less determinable and its solution shifted toward higher values. For such high photobleaching rates, including photobleaching explicitly in the probe simulations will most likely improve FISIK’s ability to accurately estimate the interaction rates.

All in all, these tests demonstrate that FISIK is able to extract molecular interaction parameters from sub-stoichiometrically labeled single-molecule imaging data, although with some systematic shifts due to the limited resolution and SNR inherent to light microscopy. For realistic SNRs commonly encountered in single-molecule imaging data, the imperfections of molecule detection and subsequently tracking might have to be compensated for by reducing the labeled fraction in order to maintain good tracking accuracy. If this labeled fraction is insufficient for interaction rate estimation (especially the association rate), then our collective results suggest that combining tracking data from multiple channels, and/or combining tracking data with dense but static oligomer distribution data (Fig 5), would provide sufficient information for FISIK to estimate accurately the unknown model parameters.

## Conclusion

In conclusion, the framework that we have developed, FISIK, is a promising approach for the inference of in situ interaction kinetics from single-molecule imaging data with sub-stoichiometric labeling. By including a labeled fraction parameter in FISIK’s modeling component, and then fitting the model to single-molecule data to determine the unknown model parameters, including the labeled fraction alongside the association and dissociation rates, FISIK tackles the issue of sub-stoichiometric labeling in a generic manner. Our proof-of-principle tests demonstrate that FISIK can indeed estimate molecular association and dissociation rates using single-molecule data with labeled fractions as low as ∼0.2 and ∼0.06, respectively. Prior knowledge about the molecule density allows FISIK to estimate the unknown model parameters with greater accuracy. The proof-of-principle tests also show that FISIK’s performance critically depends on the accuracy of molecule tracking. For molecular systems where accurate tracking requires a labeled fraction that is insufficient for parameter estimation, combining tracking data from multiple channels, and/or combining tracking data with dense but static oligomer distribution data, is a viable strategy to provide sufficient information for FISIK to estimate molecular association and dissociation rates. As our tests also show, the accuracy of FISIK depends on the accuracy of the model it employs to describe the biological system under study. Various lines of experimentation will most likely be needed to define an appropriate model to use within FISIK (e.g. motion types, interdependence between motion type and interactions, and maximum oligomeric state) and to constrain the model fitting problem. With this, FISIK is expected to allow us to estimate the interaction rates between molecules in their native cellular environment, taking full advantage of the rich spatial and temporal information present in live-cell single-molecule imaging data.

## Software availability

FISIK code, to perform all of the procedures and analyses described in this work, is available on github (https://github.com/kjaqaman/FISIK).

## Supporting material

The manuscript includes 1 supporting note, 5 supporting figures, 5 supporting tables and 4 supporting videos.

## Author contributions

KJ and LRO designed and implemented the framework, performed the tests and data analysis (primarily LRO), and wrote the manuscript.

## Acknowledgments

We thank R. Yirdaw for contributing toward initial implementations of the simulation code. We thank Chad Brautigam and Satwik Rajaram for critical reading of the manuscript and helpful discussions. This work was supported by funding from The Welch Foundation (I-1901), NIH/NIGMS (R35 GM119619) CPRIT (R1216), and the UT Southwestern Endowed Scholars program to KJ.

