## Supporting Material for "FISIK: Framework for the Inference of in Situ Interaction Kinetics from single-molecule imaging data"

Includes 1 supporting note, 5 supporting figures, 5 supporting tables and 4 supporting video legends.

#### Note S1: Clean-up of the u-track output to supply target data to FISIK

Occasionally, a track got merged with and split from at the same time point. In this case, merge-split pairs canceled out each other. Specifically, suppose $N_M$ merges and $N_S$ splits occurred with the same track (call it the "base track") at the same time (Fig S5). Then, min($N_M$, $N_S$) merges and splits were paired with each other and separated from the base track. Any excess merges (if $N_M > N_S$) or excess splits (if $N_S > N_M$) stayed connected to the base track. The merge-split pairing was formulated as a linear assignment problem (LAP) (1), with the cost for pairing merge *m* with split *s* calculated from their mean intensities ($I_m$ and $I_s$, respectively) and mean step sizes ($\Delta r_m$ and $\Delta r_s$, respectively):

$$pairing\ cost(m,s) = \frac{\max(I_m, I_s)}{\min(I_m, I_s)} \times \frac{\max(\Delta r_m, \Delta r_s)}{\min(\Delta r_m, \Delta r_s)}. \quad (8)$$

While this condition was overall infrequent (~0.3% of tracks), we corrected it for the sake of completeness.

### Supporting Figures

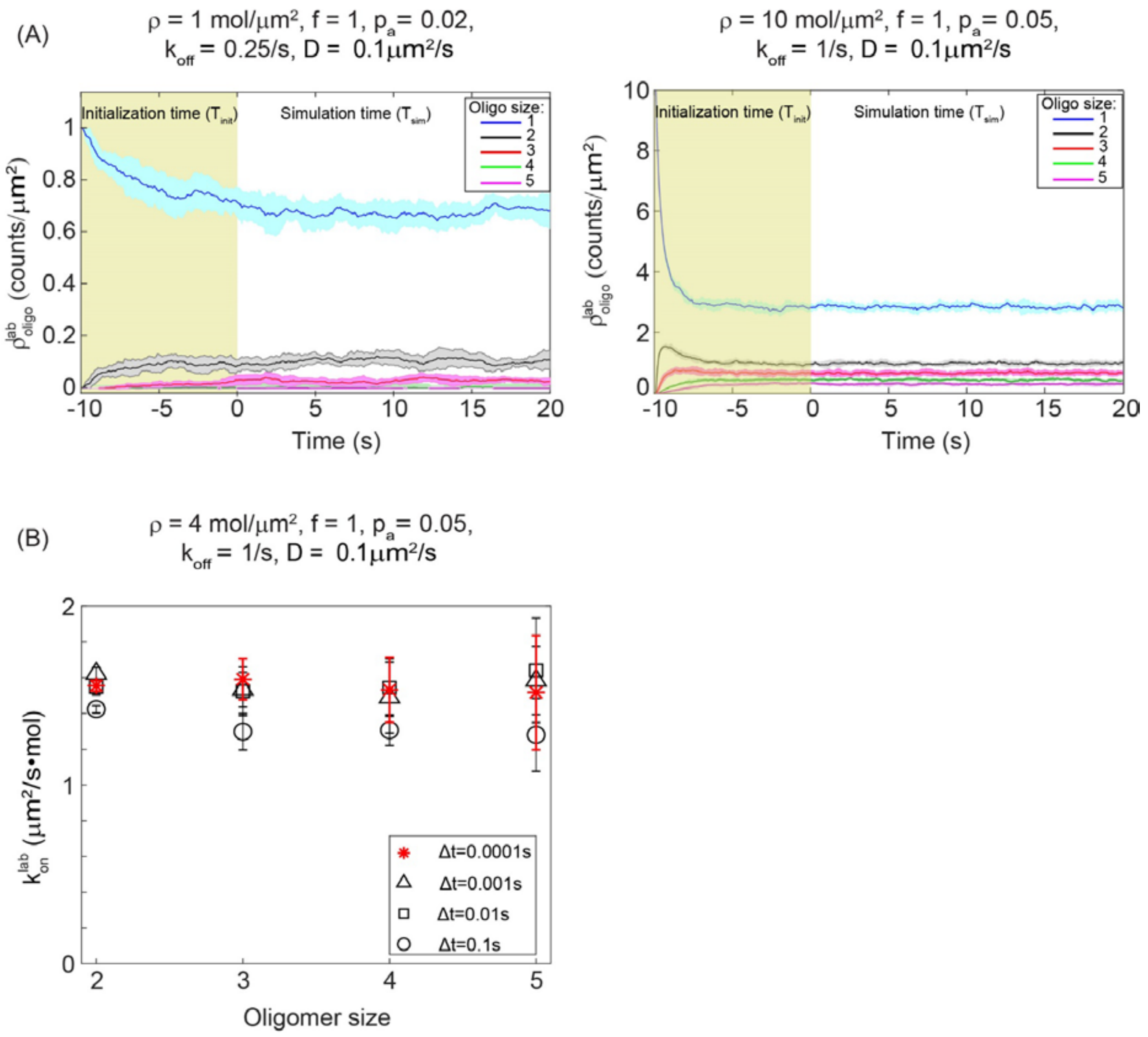


**Figure S1. Simulation strategy considerations.** (**A**) Labeled oligomer density as a function of simulation time for the indicated model parameters, shown as mean ± 1 standard deviation from 10 simulations. Yellow-green background indicates initialization time ($T_{init}$). White background indicates simulation time for output ($T_{sim}$). $T_{init}$ = 10 s is sufficient to reach steady state even for the model simulated in (A), which has one of the slowest dynamics among the simulated parameter combinations in our study. (**B**) Labeled molecule association rate as a function of the simulation time step (Δt), shown as mean ± 1 standard deviation from 10 simulations. Correcting for the finite Δt yields equivalent simulation output for Δt in the range $10^{-4}$-$10^{-2}$. Δt = 0.1 is however too large and the correction seems to not compensate for it. Note that, for analysis, all trajectories are downsampled to an output time step of 0.1, regardless of the simulation time step.

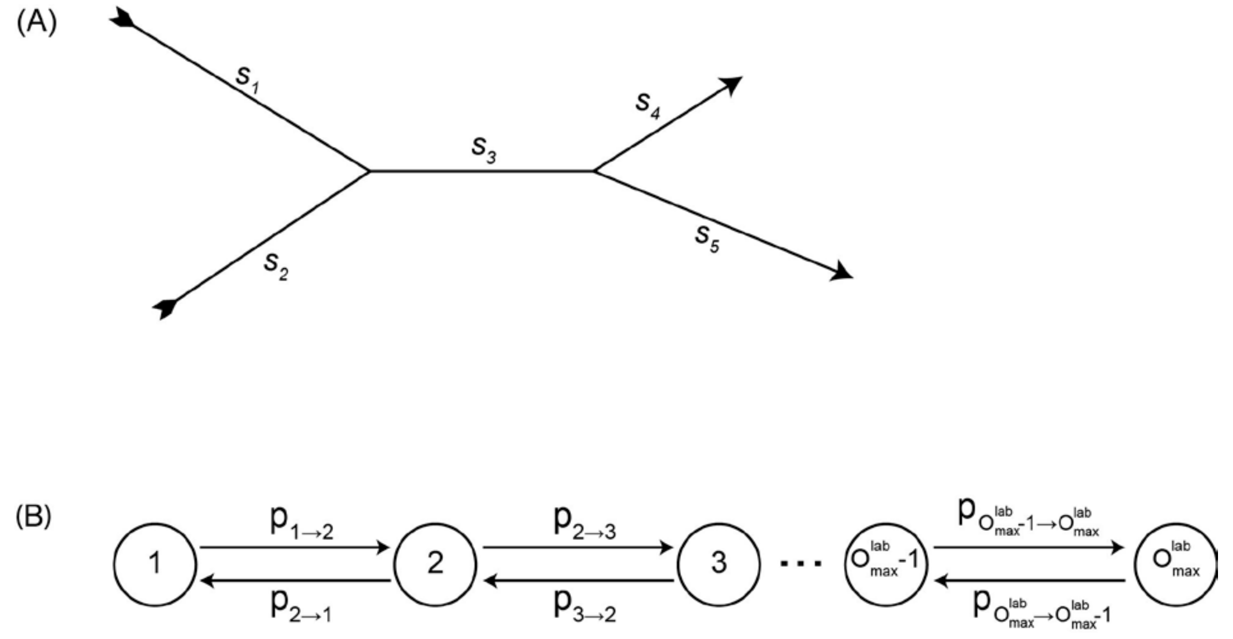


**Figure S2. Single-molecule data analysis and intermediate statistics calculation considerations.** (**A**) Example of two particles merging and splitting. For assignment of particle oligomeric state, this "compound track" (which includes information for both particles) is divided into segments $s_1$ and $s_2$ before the merge, segment $s_3$ after the merge (but before the split), and then segments $s_4$ and $s_5$ after the split. This merging and splitting history provides constraints (Eq. 3) for the objective function minimization (Eq. 2) to determine the particle oligomeric states. (**B**) Markov process representation of transitions between oligomeric states due to molecule association and dissociation (also called oligomeric sizes in the text), for calculating the labeled molecule dissociation rate (Eq. 4).

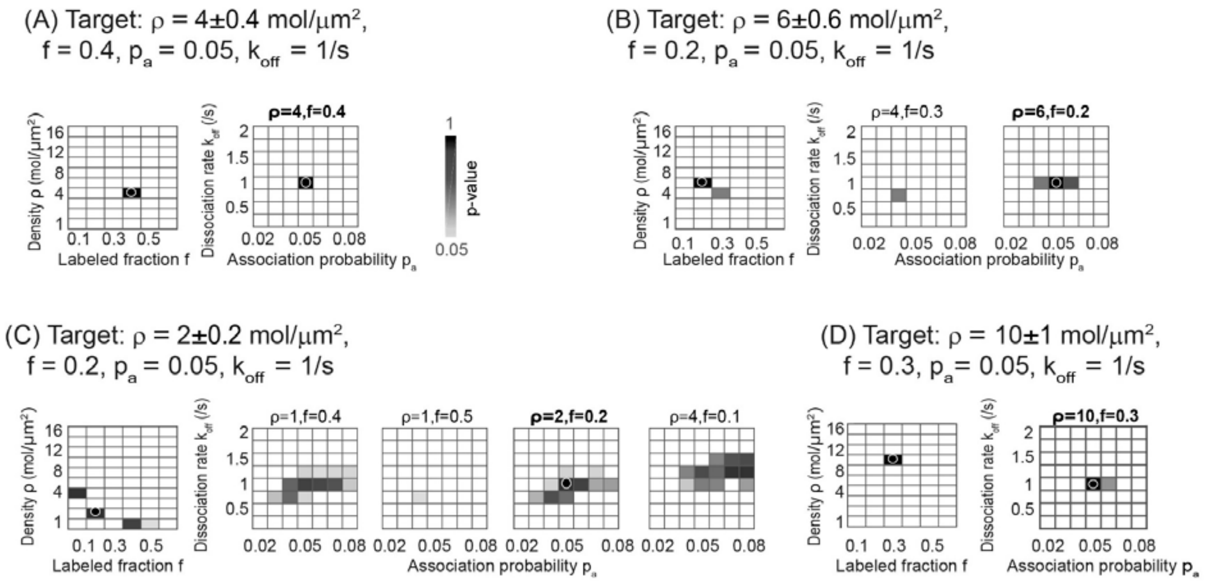


**Figure S3. Further tests of FISIK by varying molecule density and labeled fraction under conditions of relatively low density and low-medium labeled fraction (as in Fig 3).** Four dimensional p-value landscape for comparing probe trajectories to the indicated targets, as a function of the population parameters $\rho$ and $f$ and the interaction parameters $p_a$ and $k_{off}$. Detailed description as in Fig 3A. (**A**-**D**) Same as in Fig 3A, but with target $f$ increased (**A**), target ρ increased (**B**) or decreased (**C**), or both target $\rho$ and target $f$ increased (**D**). The correct probe parameters (= target parameters) had the following p-values: 0.9806 (rank 1/1 possible solutions) in (A), 0.9845 (rank 1/4 possible solutions) in (B), 0.8917 (rank 2/36 possible solutions) in (C), and 0.9999 (rank 1/2 possible solutions) in (D).

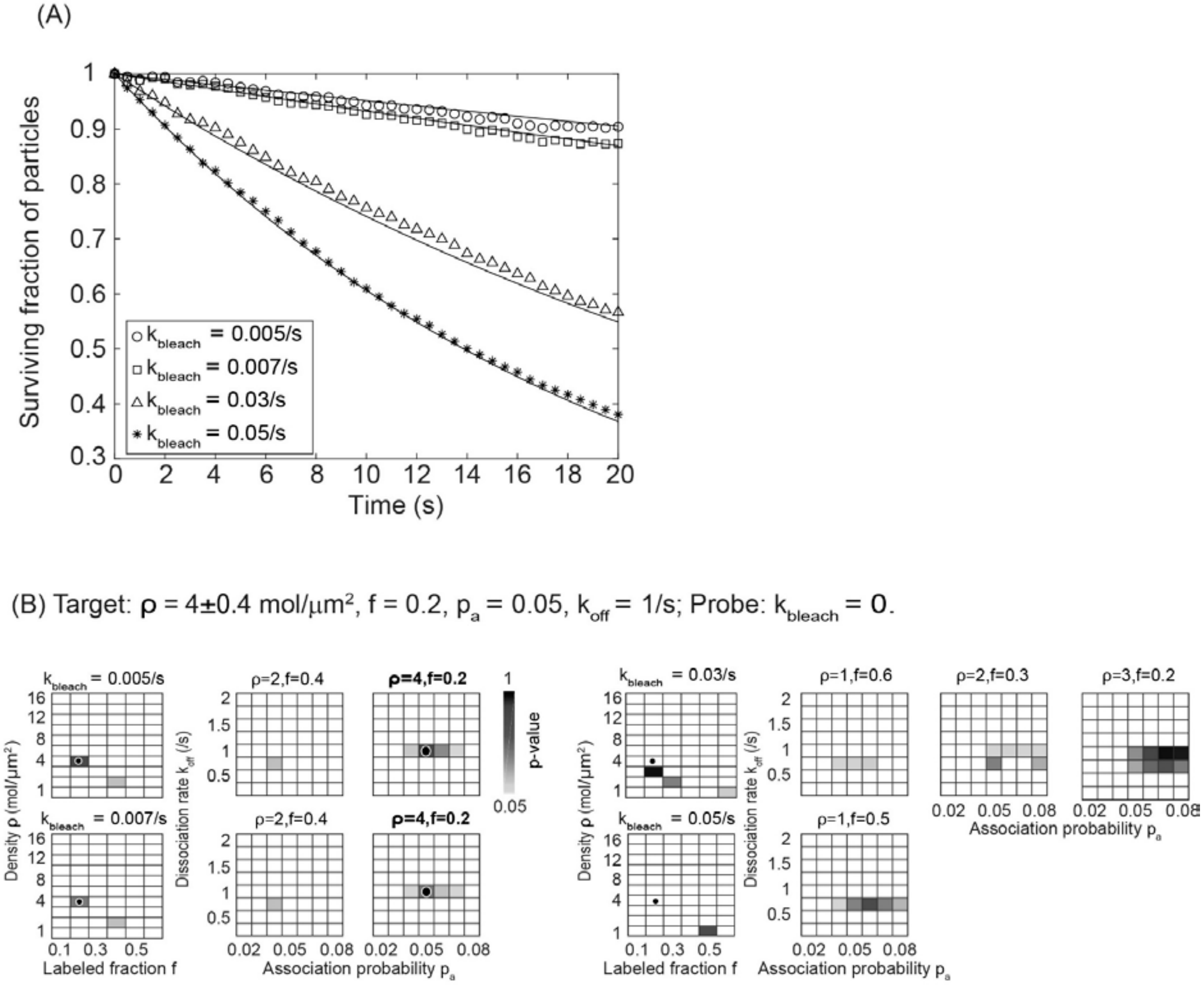


**Figure S4. The effect of photobleaching that is not accounted for on FISIK's performance. (A)** Surviving fraction of particles in target simulations where the labeled fraction decays exponentially to mimic fluorophpore photobleaching. Individual points are the measurements while solid lines are the theoretical curves for exponential decay with indicated rates. (**B**) Four dimensional p-value landscape for comparing probe trajectories to the indicated targets, as a function of the population parameters $\rho$ and $f$ and the interaction parameters $p_a$ and $k_{off}$ (total of 3360 probes). Detailed description as in Fig 3A. Targets were subject to photobleaching with the indicated rates ($k_{bleach}$), while the probes did not include any photobleaching. The correct target parameters had the following p-values: 0.9653 (rank 1/5 possible solutions) for $k_{bleach}$=0.005/s, 0.4316 (rank 1/5 possible solutions) for $k_{bleach}$=0.007/s. They were outside of the solution range for $k_{bleach}$=0.03/s and $k_{bleach}$=0.05/s.

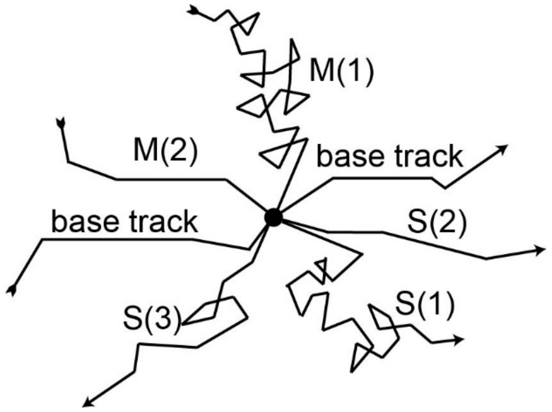


**Figure S5. Illustration of simultaneous merges and splits that get "cleaned up," as described in S1 Appendix.** In this example, 2 merges (M(1) and M(2)) and 3 splits (S(1), S(2) and S(3)) occur with the base track at the same time point. After the clean-up, two merge-split pairs (M(1)-S(1) and M(2)-S(2)) are separated from the base track, leaving only one excess split, S(3), connected to the base track.

### Supporting Tables

| $A$ (μm$^2$) | $\Delta t$ (s) | $T_{sim}$ (s) | $T_{init}$ (s) | $\Delta t_{out}$ (s) | $N_{sim}$ |
|---|---|---|---|---|---|
| 10 × 10 | 0.01 | 20 | 10 | 0.01 | 10 |
| 25 × 25 | 0.01 | 20 | 10 | 0.1 | 10 |
| 12 × 12 | 0.01 | 20 | 10 | 0.1 | 30 |
| 12 × 12 | 0.01 | 1 snapshot | 30 | N/A | 30 |

**Table S1: Simulation parameters, for probe and target simulations.** Shown are the values for simulation area ($A$), simulation time step ($\Delta t$), simulation time ($T_{sim}$), initialization time ($T_{init}$), output trajectory time step ($\Delta t_{out}$), and number of simulations ($N_{sim}$). Per the requirements of indirect inference (2), a fixed set of random number generator seeds was used in the simulations, in order to reduce the jaggedness of the objective function landscape when varying model parameters. Probe and target simulations employed different random number generator seeds. The rows refer to the following simulations: Row 1. Validation (Fig 2). Row 2. Low density (Fig 3). Row 3. High density, dynamic (Fig 4). Row 4. High density, static (Fig 5A).

| $D$ (μm$^2$/s) | $\rho$ (mol/μm$^2$) | $f$ (all rows except 3) $\phi$ (row 3) | $p_a(2\leq n\leq 5)$ | $k_{off}(n\geq 2)$ (/s) | # of parameter comb. |
|---|---|---|---|---|---|
| 0.1 | 1, 2, 3, 4, 6, 8, …, 16 | 0.1, 0.2, …, 0.6 | 0.02, 0.03, …, 0.08 | 0.25, 0.5, …, 2 | 3360 |
| 0.1 | 80,100,120 | 0.01, 0.02, …, 0.14 | 0.02, 0.03, …, 0.08 | 0.25, 0.5, …, 2 | 2352 |
| 0.1 | 80,100,120 | 0.75, 0.8, …,0.95 | 0.02, 0.03, …, 0.08 | 0.25, 0.5, …, 2 | 840 |
| 0.1 | 4 | 0.1 | 0.09, 0.10, …, 0.15 | 0.25, 0.5, …, 2 | 56 |
| 0.1 | 2 | 0.2 | 0.09, 0.10, …, 0.15 | 0.25, 0.5, …, 2 | 56 |
| 0.1 | 1 | 0.4, 0.5 | 0.09, 0.10, …, 0.15 | 0.25, 0.5, …, 2 | 112 |

**Table S2: Model parameters for generating probe landscapes.** Shown are the values for diffusion coefficient ($D$), molecule density ($\rho$), labeled fraction ($f$) or observed fraction ($\phi$), association probability ($p_a$) and dissociation rate ($k_{off}$). Note that $p_a(n > 5) = 0$. The number of probe parameter combinations is shown in the last column. Target simulations used a subset of these parameter combinations, spanning the whole range. The rows refer to the following simulations: Row 1. Low density (Fig 3). Row 2. High density, dynamic (Fig 4). Row 3. High density, static (Fig 5A). Rows 4-6. Additional probes, supplementing those in Row 1, for u-track comparison (Fig 7). These were needed because $p_a$ was shifted toward higher values when FISIK was applied to u-track targets.

| Image area (pixels$^2$) | Pixel size (nm) | PSF std (pixels) | Bit depth | Single-molecule intensity [mean, std] (a.u.) | Background mean (a.u.) | Background std (i.e. noise) (a.u.) | Mean SNR |
|---|---|---|---|---|---|---|---|
| 302x302 | 90 | 1.2 | 16 | [1000, 100] | 5000 | 140 | 7 |
| 302x302 | 90 | 1.2 | 16 | [1000, 100] | 5000 | 50 | 20 |

**Table S3: Synthetic image generation parameters.** Shown are the values for image area, pixel size, standard deviation (std) of Gaussian approximating the point spread function (PSF), image bit depth, single-molecule intensity above background, background mean, background std, and resulting signal-to-noise ratio (SNR).

| **Detection parameters** | | **SNR 7 and 20** |
|---|---|---|
| Gaussian standard deviation (pixels) | | 1.2 |
| Local maxima detection | α-value for comparison with local background | 0.1 |
| Gaussian fitting at local maxima | Do iterative Gaussian mixture-model fitting | Yes |
| | α-value for residuals test | 0.2 |
| | α-value for amplitude test | 1 |
| | α-value for distance test | 1 |

**Table S4: U-track detection parameters (Gaussian Mixture-Model Fitting).** Shown are the parameters with non-default values (based on u-track Version 2.2.1).

| Tracking parameters | | SNR 7 and 20 |
|---|---|---|
| Overall | Maximum gap to close (frames) | 2 |
| | Do segment merging | Yes |
| | Do segment splitting | Yes |
| Frame-to-frame linking cost function | Brownian search radius lower bound | 7 |
| | Brownian search radius upper bound | 7 |
| Gap closing, merging and splitting cost function | Brownian search radius lower bound | 7 |
| | Brownian search radius upper bound | 7 |
| | Merging & splitting: Minimum intensity ratio | 0.75 |
| | Merging & splitting: Maximum intensity ratio | 1.5 |
| | Merging & splitting: For an end/start, the possibility of merging/splitting is allowed only if there is no possibility of gap closing | Yes (see legend for more information) |
| | Birth and death cost | Derived from gap closing and merging/splitting cost formulae |

**Table S5: U-track tracking parameters.** Shown are the parameters with non-default values (based on u-track Version 2.2.1). As explained in the original publication (3), u-track uses particle movement and intensity information to decide on merging and splitting events. After the original publication, a new constraint was added to help further distinguish between specific interactions and particles passing by each other. Specifically, merging and splitting events can be limited to ends and starts that do not have the possibility of gap closing, a constraint used in our work here.

#### Supporting video legends

**Video S1. Synthetic time-lapse (left) and detected objects (right), with an average SNR = 7.** Time-lapse corresponds to a target simulation with parameters $\rho$ = 2 ± 0.2 mol/µm$^2$, $f$ = 0.2, $p_a$ = 0.05, $k_{off}$ = 1/s and $D$ = 0.1 µm$^2$/s. Frame rate = 10 Hz. Image size = 27 µm × 27 µm. See text for image generation details.

**Video S2. Particle tracking in time-lapse with an average SNR = 7.** Time-lapse same as in Video S1. Trajectories are shown as dragtails of length 15 frames. Merges are shown as yellow diamonds, splits as green diamonds, and closed gaps as cyan circle.

**Video S3. Synthetic time-lapse (left) and detected objects (right), with an average SNR = 20.** Time-lapse corresponds to a target simulation with parameters $\rho = 2 \pm 0.2$ mol/µm$^2$, $f = 0.2$, $p_a = 0.05$, $k_{off} = 1$/s and $D = 0.1$ µm$^2$/s. Frame rate = 10 Hz. Image size = 27 µm × 27 µm. See text for image generation details.

**Video S4. Particle tracking in time-lapse with an average SNR = 20.** Time-lapse same as in Video S3. Trajectories are shown as dragtails of length 15 frames. Merges are shown as yellow diamonds, splits as green diamonds, and closed gaps as cyan circle.
